# The impact of sex on the immune system explored at the single-cell level

**DOI:** 10.1101/2025.07.27.667062

**Authors:** Seyhan Yazar, Jose Alquicira-Hernandez, Kristof Wing, Anne Senabouth, Stacey Andersen, Kirsten A. Fairfax, Alex W. Hewitt, Joseph E. Powell, Sara Ballouz

## Abstract

Sex has a key role in disease susceptibility, in particular autoimmunity. Sex differences of the immune system stems from genes and their interactions with intrinsic and external factors. However, the cellular-level factors influencing sexual dimorphism is not fully understood. We thus examined immune sex differences at single-cell resolution to dissect the genetic impacts. Female-biased sex-differentially expressed genes (DEGs) in multiple immune cells were involved in TNFɑ signalling, whereas male DEGs were enriched for ribosomal-related functions. While cis-eQTLs were less common on sex chromosomes, we identified over 1000 sex-specific eQTLs and 51 sex-interacting eQTLs on autosomes. When we examined the effect of genetic control on sex-DEGs, we found genetic variants affecting the female-biased expression *FCGR3A* in NK cells (rs2099684) and *ITGB2* in Monocytes (rs760462), both of which are associated with systemic lupus erythematosus. Our work reveals novel biases masked in bulk analyses and highlights sexually dimorphic genes and pathways at baseline.

## Main Text

Sex differences in the immune system has a critical role in autoimmune disease susceptibility, progression and outcome. These differences are evident in immune parameters such as antigen presentation strength, duration and intensity of immune responses, all of which vary between females and males. For instance, in females, vaccine responses are typically stronger, and the rates of chronic viral infections and degree of viraemia (e.g., in HIV) are lower. However, the disadvantage of these robust immune responses is that they predispose females to higher rates of autoimmunity and inflammatory diseases. In contrast, males are more likely to develop non-reproductive cancers and become affected by bacterial and parasitic infections than females(Klein and Flanagan, 2016b; Wilkinson et al., 2022).

At the cellular level, these clinical differences are quantifiable between males and females in both the innate and adaptive immune compartments. While males exhibit a greater number of circulating natural killer (NK) cells, females display a higher frequency of B cells(Abdullah et al., 2012). Additionally, females show heightened T cell cytotoxic and inflammatory responses, particularly following multiple stimulations(Hewagama et al., 2009). At the gene expression level, most sexually dimorphic traits arise on and from the sex chromosomes. The X chromosome uniquely encodes several critical immune-related genes, including interleukin receptors (*IL2Rγ*; *CD132*), chemokines (*CXCR3*), toll-like receptors (*TLR7*, *TLR8*), and genes involved in T-cell and B-cell effector functions (*BTK*, *IKKγ*, *NFκBRF*)(Fish, 2008), as well as regulatory molecules such as *FOXP3* and *CD40L* (CD154). Their differential expression is a potential driver of sex-specific immune phenotypic variation and disease.

While differences are observable at the phenotypic and cellular levels as described, they are not apparent at the genetic level. Despite the identification of over 2,000 single nucleotide variations (SNVs) associated with autoimmune diseases, there remains a limited understanding of sex-differentiated genetic architecture. Specifically, few of these SNVs demonstrate sex-specific effects. For example, polymorphisms of *TLR7* located on the X-chromosome are a well-characterized risk factor for the autoimmune condition Systemic Lupus Erythematosus (SLE) which has a 9:1 prevalence in women compared to men(Yacoub Wasef, 2004). We propose that the genetic regulation of sex-biased gene expression may provide additional evidence to clarify this unresolved research area.

Characterisation of sexual dimorphism in the adaptive and innate immune systems has previously focused on investigation of a-priori defined subsets of immune cells or used bulk analyses. This hypothesis-driven research has demonstrated key phenotypic differences between male and female immune systems, including in cell counts, population dynamics, and cytokine production, however, may result in biased and insensitive analyses. As sex differences in the immune system arise from cellular diversity, cell population composition, and cellular activity(Klein and Flanagan, 2016a), the application of single-cell analyses uniquely permits the unbiased characterisation of sex differences on the peripheral immune system.

Herein, we combine the power of single-cell gene expression (sc-RNA-seq) and genetic variation to assess sex differences at the cellular level in peripheral blood mononuclear cells (PBMCs) in a large cohort (**Fig. 1A**). We evaluated cell-type proportions, sex-biased gene expression and sex-specific and sex-interacting estimated quantitative trait loci (eQTLs) along with co-expression networks (**Fig. 1B**). By conditioning our analyses on sex and/or cell type, we disentangle the contributions of each to further understand the causes and consequences of sexual dimorphism in circulating immune cells.

**Fig. 1:**
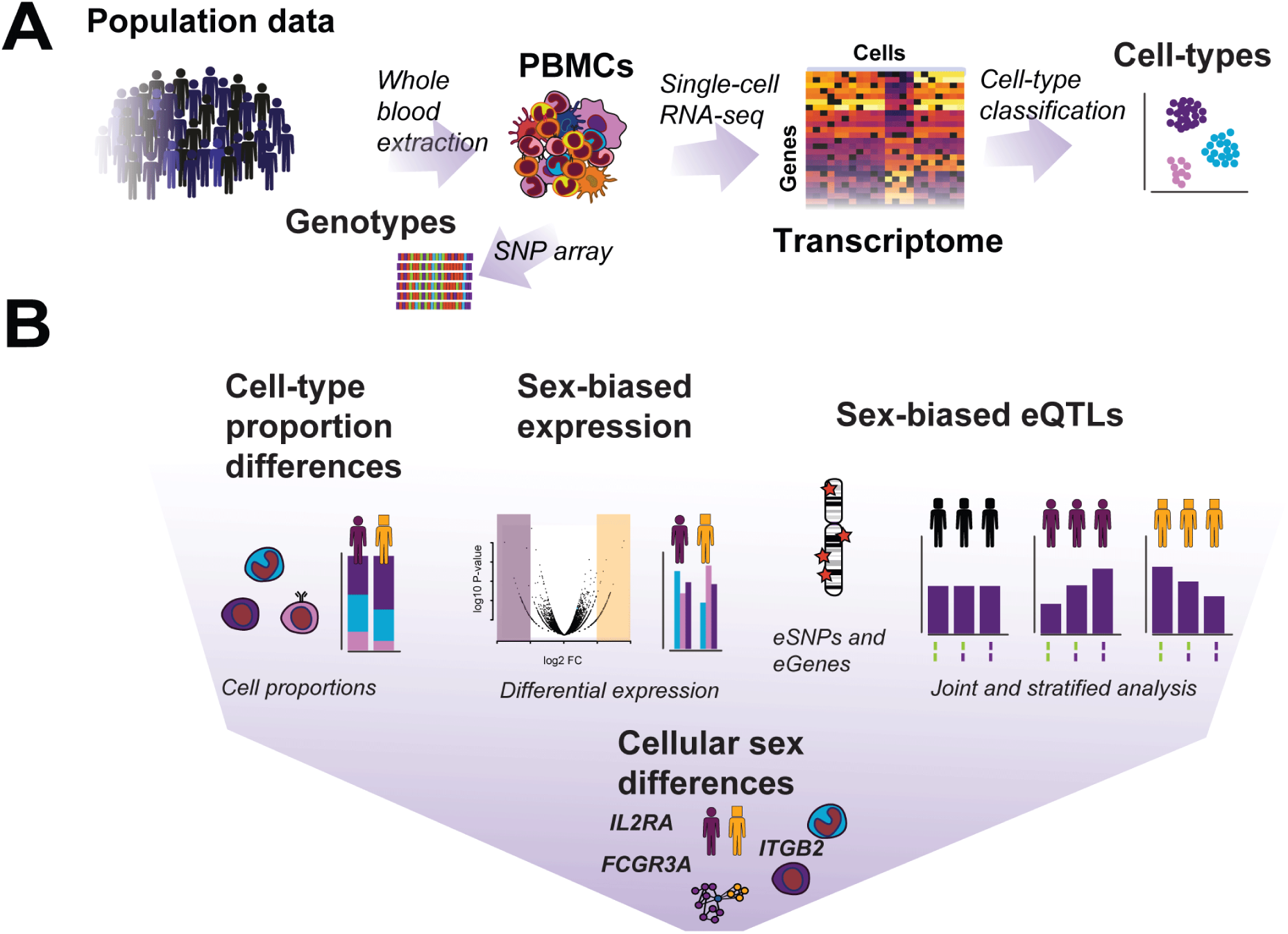
Overview of study. **(A)** Cohort: OneK1K of 982 individuals, with 564 females and 417 males. After single-cell sequencing of ∼1000 cells per individual, 1,267,758 PBMCs were classified into 30 cell-types. (B) Study design. We assessed cell-type proportion differences, sex-biased expression through differential expression, and sex-biased eQTLs through a statistical association framework.

## Results

### Single-cell data reveals novel cell-type proportion differences

We classified ∼1.25 million cells from 982 individuals from the OneK1K study (Yazar et al., 2022) (565 females, 418 males) into 30 transcriptionally distinct cell types using the Azimuth classification tool (Hao et al., 2021b) (**Fig. 2A**). Calculating proportions of each cell type per individual, we observed clear compositional differences between the sexes (**Fig. 2D-E**). In males, we found higher proportions of CD14+ Monocytes, Dendritic Cells (DCs), Natural Killer Cells (NK), NK Proliferating, CD8+ Proliferating, T-effector memory (TEM), and T-central memory (TCM) cells. In females, we found significantly higher proportions of B cells, CD4+ Naive T cells, NK CD56+, and regulatory T cells (Tregs). Most of these differences have been reported in the literature(Abdullah et al., 2012; Al-Attar et al., 2016; Huang et al., 2021; Li et al., 2019; Robinson et al., 2022b), yet some were novel including Tregs and DCs.

**Fig. 2:**
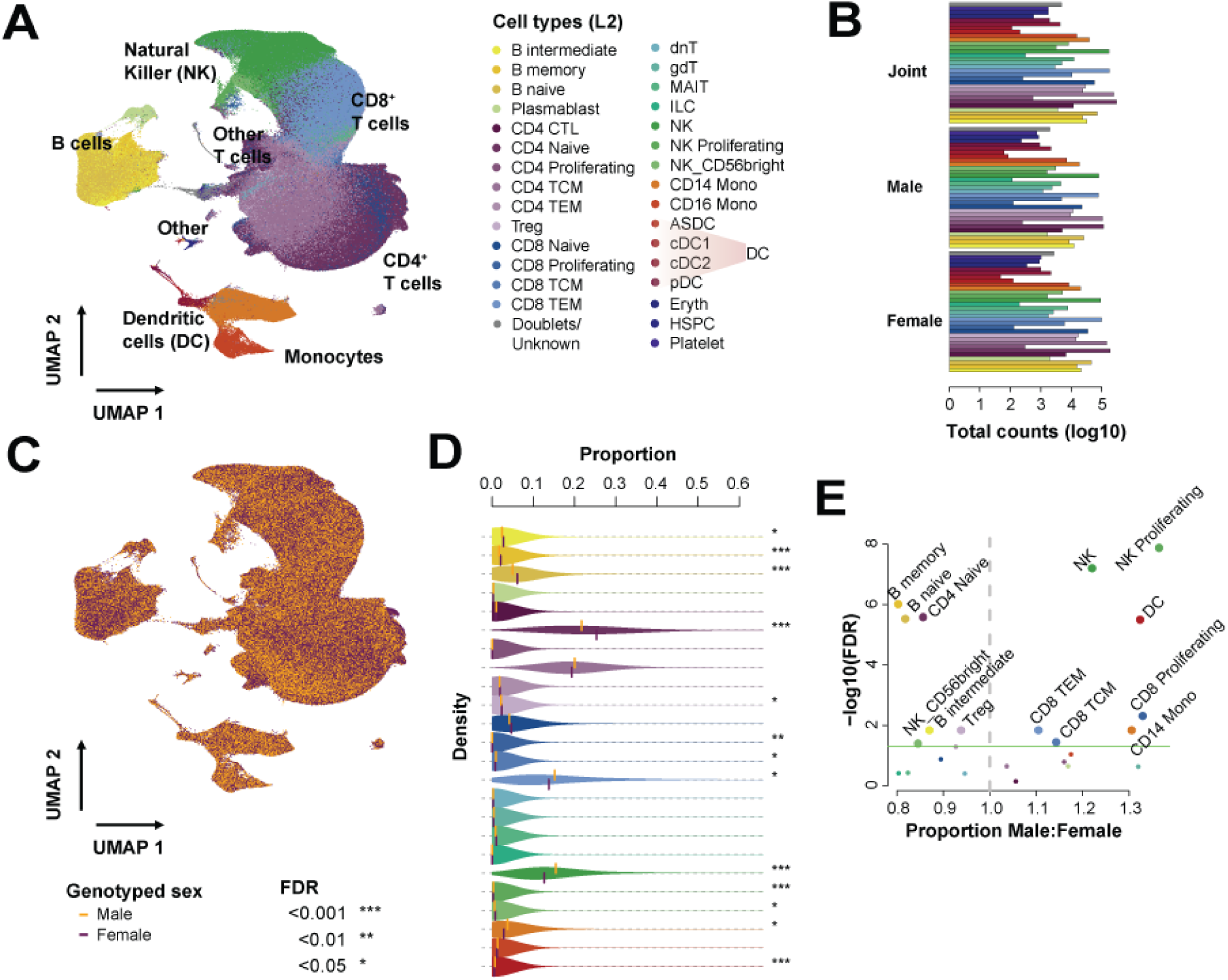
Distributions of cell-type proportions across sex. (A) UMAP of all 1,267,758 cells, coloured by cell type. Dendritic cell labels (ASDC, cDC1, cDC2 and pDC) were combined for all downstream work. (B) Total cell counts in data, for all individuals, then split by sex. (C) UMAP of cells coloured by sex. (D) Distribution of proportions of each cell-type. Females are shown on the top half of violin plot, and males bottom half. Significance of differences based on a two-sided Wilcoxon-test are indicated by the asterisk (* FDR<0.05, ** FDR<0.01, *** FDR<0.001). (E) Male to female ratios versus FDR based on a F-test (proportions)

In the innate immune system, CD14+ Monocytes were present in higher proportions in males (3.71% in males versus 2.84% in females, FDR∼0.014). Higher proportions of monocytes have been reported in male infants(Bellamy et al., 2000) and some ethnicities (Chen et al., 2016), but is not a well-established phenomenon between the sexes(Jiang et al., 2014; Puissant-Lubrano et al., 2018). No proportional difference was present in CD16+ Monocytes, consistent with others’ observations(Puissant-Lubrano et al., 2018). Due to low overall counts, we combined the DC subtypes: Plasmacytoid (pDC), conventional cell 1 (cDC1), conventional cell 2 (cDC2), and *AXL*+ DC (aDC) into a broad dendritic cell category. We observed a strong relationship between sex and DC proportion, with higher percentages in males (0.64% in males vs 0.48% in females, FDR<0.001), this has not been previously observed in flow-cytometric studies of immune variation.

In the adaptive immune compartment, we found higher proportions of Tregs in our female samples (2.18% in males vs 2.32% in females, FDR∼0.0145), where the opposite or no difference was typically reported (Kverneland et al., 2016; Robinson et al., 2022b). Tregs vary during the female menstrual cycle in response to oestrogen (Arruvito et al., 2007) and other sex steroids, so these results are potentially confounded with sex hormone levels. As we did not have associated hormone related information, we used age as a proxy to test our hypothesis. As an in-depth analysis with age is beyond the scope of this work (Sopena-Rios et al., 2024), we performed a simple linear regression analysis (**Supplementary Fig. S1, Supplementary Table S2**) and identified no significant correlation with Tregs with age (Churov et al., 2020).

### Sex-differential expression by cell type reveals novel sex biases obscured in bulk analyses

To measure differences in response and activity across the transcriptome, we performed a differential expression analysis between the sexes to identify sex-biased gene expression across cell types. We detected a total of 101 differentially expressed genes (sex differentially expressed genes (DEGs)) across 24 cell types: 23 genes with male-biased expression and 81 genes with female-biased expression (**Fig. 3A, Supplementary Table S3-4**). Many of these genes (62%) were unique per cell type (**Fig. 3B**), and the most recurrent were known sex-specific marker genes (*i.e.*, *XIST* and *RPS4Y1*). Within the set of shared genes, three (*CD79A*, *TSC22D3* and *JUN*) varied in their direction of sex bias across the cell types. These genes showed female-biased expression in B-cells but were more highly expressed in male dendritic cells. Of the three, only *TSC22D3*, an anti-inflammatory protein glucocorticoid (GC)-induced leucine zipper (also known as GILZ), is a known sex-biased and X-linked gene, while the others were novel to our work. Important to note that of the 101 genes, only 21 of these genes were on the sex chromosomes (16 on the X and 5 on the Y), implying additional sex-specific regulation of autosomal genes.

**Fig. 3:**
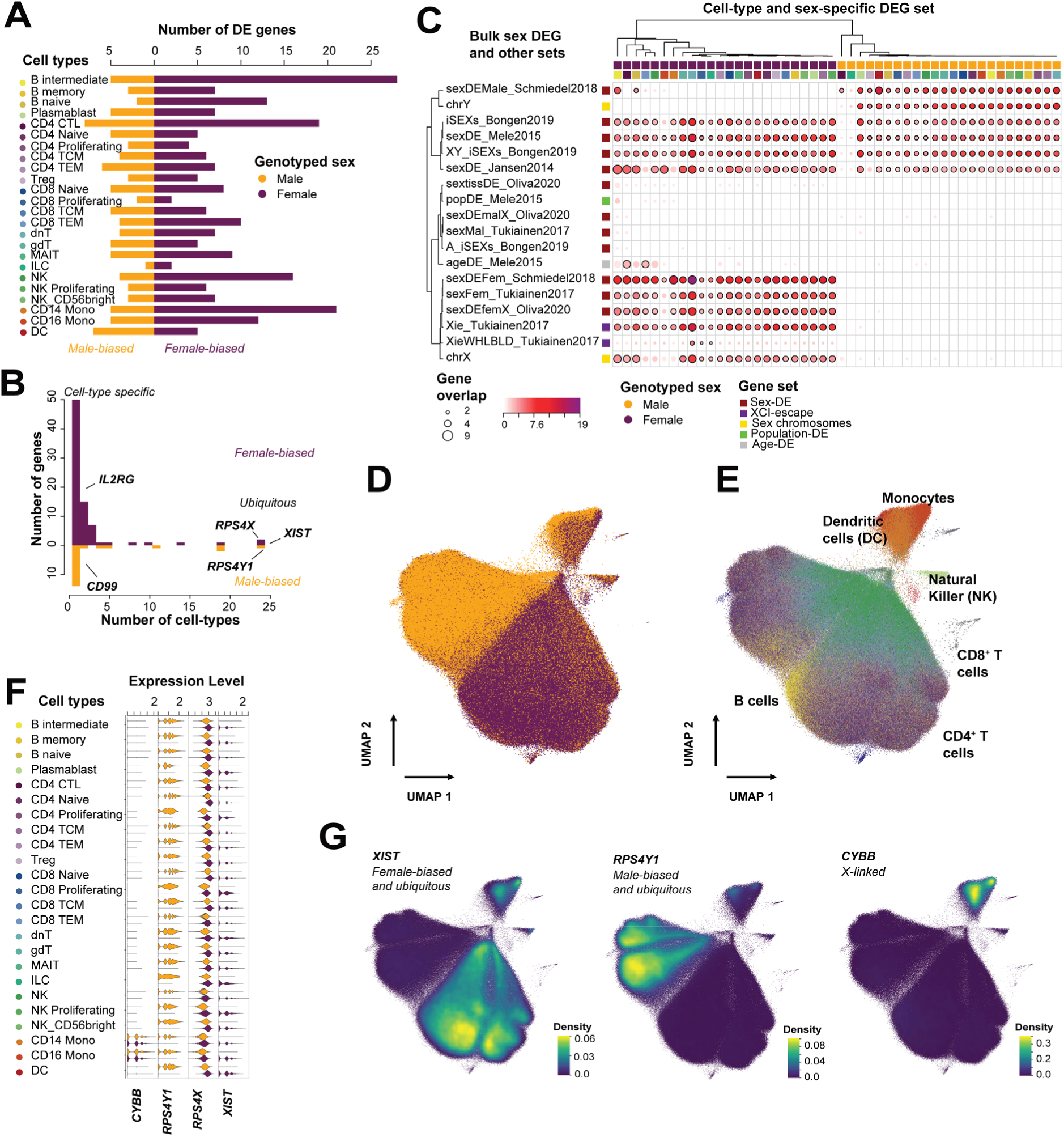
Sex-differential expression. (A) Number of sex-differentially expressed genes using the Wilcoxon-test per cell-type (two-sided). (B) Recurrence of these DEGs shows mostly unique genes, but sex-specific markers recurrent across all cell-types (ubiquitous). (C) Gene set enrichment analysis of DEGs showing enrichment of sex-specific gene sets. Additionally, age related enrichment in DEGs of CD8 TEM and CD4 CTL, and population related genes in B intermediate cells. (D) Exploiting variation of the genes on the XY chromosomes to visualize sex through a UMAP. (E) Then coloured by cell-types showing that some cells-types are partially encoded on the sex chromosomes. (F) Violin plots and (G) UMAPs of cell-type specific markers (*XIST* and *RPS4Y1*) with clear sex differences. Expression of X-linked genes *CYBB*, with no sex-bias and *RPS4X* which is differentially expressed (violin plot only)

We tested for overlap of our results with known sex DEGs from bulk RNA-seq studies and X-linked gene sets of interest (Bongen et al., 2019; Jansen et al., 2014; Melé et al., 2015; Oliva et al., 2020; Schmiedel et al., 2018; Tukiainen et al., 2017) (**Fig. 3C, see Methods**). Of the 101 genes, 51 have been identified as sex DEGs in bulk studies of whole blood or other tissues. We also noted consistent sex-biased expression profiles. That is, if it is up-regulated in females (female-biased), then this expression pattern is maintained at cellular resolution (**Supplementary Table S4**). The exceptions include *FLNA* (filamin A) and *SAT1* (Spermidine); these are male-biased sex-DEGs from bulk, but both appear female-biased in our data. This inversion of biased expression could also be due to bulk analyses masking expression or could be linked to variation in X-escape of these genes in different tissues and cell types. Many of the 50 novel genes were unique to a cell-type, highlighting how improving cellular resolution through single-cell analysis can extract signals obscured in bulk. Of these, 7 genes were cell-type specific in males, and 24 genes in females. We also identified novel recurrent sex-biased genes – i.e., those that appear in more than one cell-type. In males, the most recurrent autosomal gene across the cell types was *RPLP1* (Ribosomal Protein Lateral Stalk Subunit P1), a component of the 60S ribosomal subunit (12/24 cell types). Other ribosomal proteins that appeared as sex DEGs (*RPL17*, *RPS29, RPL36A, RPL37A* and *RPS15A*) were also upregulated in males. Upregulation of ribosomal components are linked to cell growth and proliferation, but also cancer (Guimaraes and Zavolan, 2016).

### Immune pathways are enriched in sex-biased genes

To quantify the overlap of the sex-biased genes with function, we tested for gene set enrichment of pathways and gene groups (**Supplementary Fig. S2**). The female-biased sex-DEGs in B intermediate, B naive, CD14+ Monocytes, CD8+ TEM and NK cells were enriched for the TNFɑ (Tumor necrosis factor alpha) signalling pathway (adjusted p: B intermediate ∼2.04×10^-9^, B memory ∼6.74×10^-3^, CD14+ Mono ∼2.81×10^-2^, CD8+ TEM ∼2.94×10^-3^, and NK ∼3.41×10^-4^), which included genes regulated by NF-kB in response to TNFɑ expression. TNFɑ is a cytokine used by the immune system for cell signalling, and dysregulation of NF-kb has been linked to inflammatory and autoimmune diseases (Barnabei et al., 2021). The genes driving this enrichment were similar for most cell types and included *JUN*, *DUSP1*, *DUSP2*, *IER2*, *ZFP36, CD69, CD83, SAT1, KLF6,* and *PPP1R15A*. Many of these genes were also involved in the hypoxia pathway that was significantly enriched in B cells (p-adjusted: B intermediate ∼9.72×10^-5^, B naive ∼ 5.73×10^-3^). Furthermore, these genes were involved in cellular proliferation, specifically of T-cells (Grosche et al., 2020). Interestingly, the CD14+ Monocytes had a distinct set of genes driving the TNFɑ enrichment, including *G0S2*, *NFKBIA*, and *PLAUR*. These genes were also involved in the inflammatory response (CD14+ Monocytes p-adjusted ∼8.88×10^-4^), along with *CD14* and *EMP3,* both linked to monocyte differentiation/proliferation. In CD4+ CTLs, the interferon-gamma response (p-adjusted∼3.77×10^-9^) and allograft rejection (p-adjusted∼2.20×10^-2^) were enriched in female-biased sex DEGs. Some genes were shared between these two pathways, including *CD2, GZMA, HLA-A, HLA-E, IL2RG, FLNA,* and *CCL5*. The sex-specificity of all these genes is unclear as they have broad functions, but this could be due to the higher activity of these pathways in females. Monocytes are reported to have increased functional activity in females, summarized as primed IFN/immune pathways and overexpression of immune genes at basal levels(So et al., 2021). Male-biased sex-DEGs were enriched for ribosomal-related functions and pathways. These include rRNA processing, translation, and metabolism of RNA pathways (**Supplementary Fig. S2B**).

One mechanism for sex differences in gene expression is believed to occur through gene regulation by sex hormones and their receptors in immune cells (Bhatia et al., 2014). To evaluate this hypothesis, we tested for enrichment of sex hormone receptor target gene sets (oestrogen receptors *ESR1*, *ESR2*, and androgen receptor *AR*) from MotifMap (Daily et al., 2011). We found no enrichment of the receptors target genes in the differentially expressed genes (**Supplementary Fig. S2F**). Instead, genes with female-biased expression in CD14+ Monocytes were enriched for the *ESR1* TF targets gene set from ENCODE (*G0S2*, *ITGB2, MALAT1*, *MT2A*, *MYL6*, *NFKBIA*, *S100A10*, and *S100A11*), as were CD16+ Monocytes (*ANXA1, MALAT1,* and *TUBA1A*). Additionally, *LGALS2* (sex-DE in CD14+ Monocytes) was a target of *ESR2* (ERꞵ). Monocyte cell counts decrease as oestrogen levels increase through the Fas/FasL system and the ERꞵ receptor (*ESR2*) (Mor et al., 2003). Thus, our analysis highlights the role of oestrogen in differing monocyte cell counts and their functional activity.

### Genes responsible for establishing immune cell type identity are mostly distinct from those determining sex identity

Cell-type transcriptional identity is described by the expression levels of marker genes, with a few of these genes located on the sex chromosomes. These genes are not merely markers but potentially also functional elements. We examined intersection between the sex chromosome genes, sex-DEGs (including autosomal) and known cell-type marker genes, to assess their impact on cellular and sex-linked identity. We first evaluated the variation of sex-chromosomes genes within a dimension reduction framework (UMAP), which allowed us to visualize the influence of these genes on cell identity agnostic of their sex-biased expression profiles. Using the variation of genes expressed on the X and Y chromosomes, we detected a clear separation by sex (**Fig. 3D**) and a strong clustering by the myeloid and lymphoid lineages, with monocytes and dendritic cells split from the remaining cells (**Fig. 3E**). This separation indicates an important role for a subset of sex chromosome genes in monocyte and dendritic cell identity.

To further explore this finding and focus in on a potential set of sexually dimorphic and immune related genes, we tested for overlap of the sex-biased genes (sex DEGs) and cell-type marker genes. For this, we utilised the L2 cell-type classification marker genes from the Azimuth reference panel. Overall, of the 215 cell-type marker genes, 21 were sex-biased genes. When we looked at the cell type specific distribution of these 21 genes, we found 16 of them were both cell-type markers and sex DEGs within the same cell type. These included *CD79A* in B intermediate and naïve cells, *CXCR4* in B naïve cells, *CD14*, *G0S2* and *S100A8* in CD14+ monocytes, *FCGR3A* in CD16+ monocytes, *FGFBP2*, *GZMA*, *GZMH* in CD4 CTLs, *GZMK* in CD4 TEM, CD8 TEM, and MAIT cells, *KLRC1* in gDT, *NKG7* in MAIT and finally *FCER1G* in NK cells. Of these, none were on the sex chromosomes.

We then repeated the differential expression analysis between cell types (one vs all) and expanded the marker gene list to test their overlap with the sex DEGs. We identified a total of 1,749 marker genes across the cell types (FDR <0.05, |log2FC| > 0.25), with 55 (3%, FDR∼1) of these genes located on the X chromosome (**Supplementary Table S5**). The distribution of the marker genes across all the chromosomes shows no deviation from the null (𝟀𝟀^2^ p-value = 0.25), except with enrichment for genes on chromosome 19 (160 genes, FDR∼0.02) and mitochondrial genes (8 genes, FDR < 0.001). Overall, 53 marker-like genes overlapped with a sex-biased gene within that same cell-type. The overlaps range from 0% (e.g., Tregs and Plasmablasts) to 61.54% (CD14+ Monocytes, FDR∼4.84×10-6), with 17% overlap on average (**Supplementary Table S7**). Yet despite 24 of the 55 X chromosome marker-like genes showing sex-biased expression, only 5 were within the same cell-type. These included *EIF2S3* (CD4 Naive), *PLP2* (DC), *RPS4X* (CD4+ and CD8+ Naive), *SMC1A* (NK proliferating) and *TSC22D3* (B intermediate and B naive). Remarkably, one of the strongest markers for the myeloid lineage, *CYBB*, an X-linked gene, is not differentially expressed by sex (**Fig. 3G**).

The remaining 49 autosomal sex-biased and marker genes could potentially play crucial roles in cell function and sexual dimorphism of immune traits. For example, the autosomal gene *CCL5* that is a marker for CD8+ T cells, shows cell-type specific expression and is differentially expressed by sex, with higher expression in females in CD8+ TCM cells (log2FC=-0.22, FDR∼2.02×10^-22^). Increased expression of *CCL5* is potentially linked to improved anti-viral immunity through lymph-node and splenic homing of viral-specific CD8 T cells.

### Sex-specific cis-eQTLs at the single-cell level are mostly cell type specific

Sexually dimorphic phenotypes may partly derive from genetic effects and their interactions with the environment. Several eQTLs have been identified to show sex-biased or sex-interacting effects using bulk RNA sequencing(Dimas et al., 2012; Kukurba et al., 2016; Oliva et al., 2020) and more recently using single cell sequencing of lymphoblastoid cell lines(Jones et al., 2024) and PBMCs from Asian Immune Diversity Atlas(Tomofuji et al., 2024). To understand the impact of sex on genetic control of gene expression at the single cell level at the population scale, we tested for cis-eQTLs on the autosomes (**Fig. 4**) and sex chromosomes (**Fig. 5**) both jointly and stratifying by sex (**Fig. 4A, Supplementary Table S9**). We found 14,488 autosomal eQTLs across 21 cell types in our joint analysis (**Fig. 4B, Supplementary Table S10**). In our stratified analyses, we found fewer eQTLs overall, which was likely due to reduced power in the stratification and stringent filtering, with 1,041 eQTLs in females and 998 eQTLs in males (**Fig. 4B, Supplementary Table S11 & S12**). Between cell types, many significant eQTLs were unique *i.e.,* cell type specific (**Fig. 4C**). A total of 959 eGenes were observed in males, with 39 (4%) occurring in more than two cell types. In females, of the total of 991 eGenes, 42 were in two cell types, and four genes were in three (4.6% >1). Additionally, 14 female-specific eGenes were in the MHC region, compared to 16 male-specific eGenes. We compared the Rho estimates between the sexes in the novel associations (**Fig. 4D**) and noticed that most of these eQTLs had smaller estimates than joint eQTLs on average (mean |Rho|∼0.3). This reflects low but significant effects of SNPs on sex specific eGenes. Although the SNPs differed, 128 eGenes were found to be common in both male and female novel associations. The majority of these eGenes (113) showed significance in distinct cell types, indicating that a small subset of genes may have sex-specific and cell type-specific regulatory mechanism. The 15 eGenes in the same cell types had variants that were not in linkage disequilibrium, suggesting these eGenes were also under different genetic regulatory control.

**Fig. 4:**
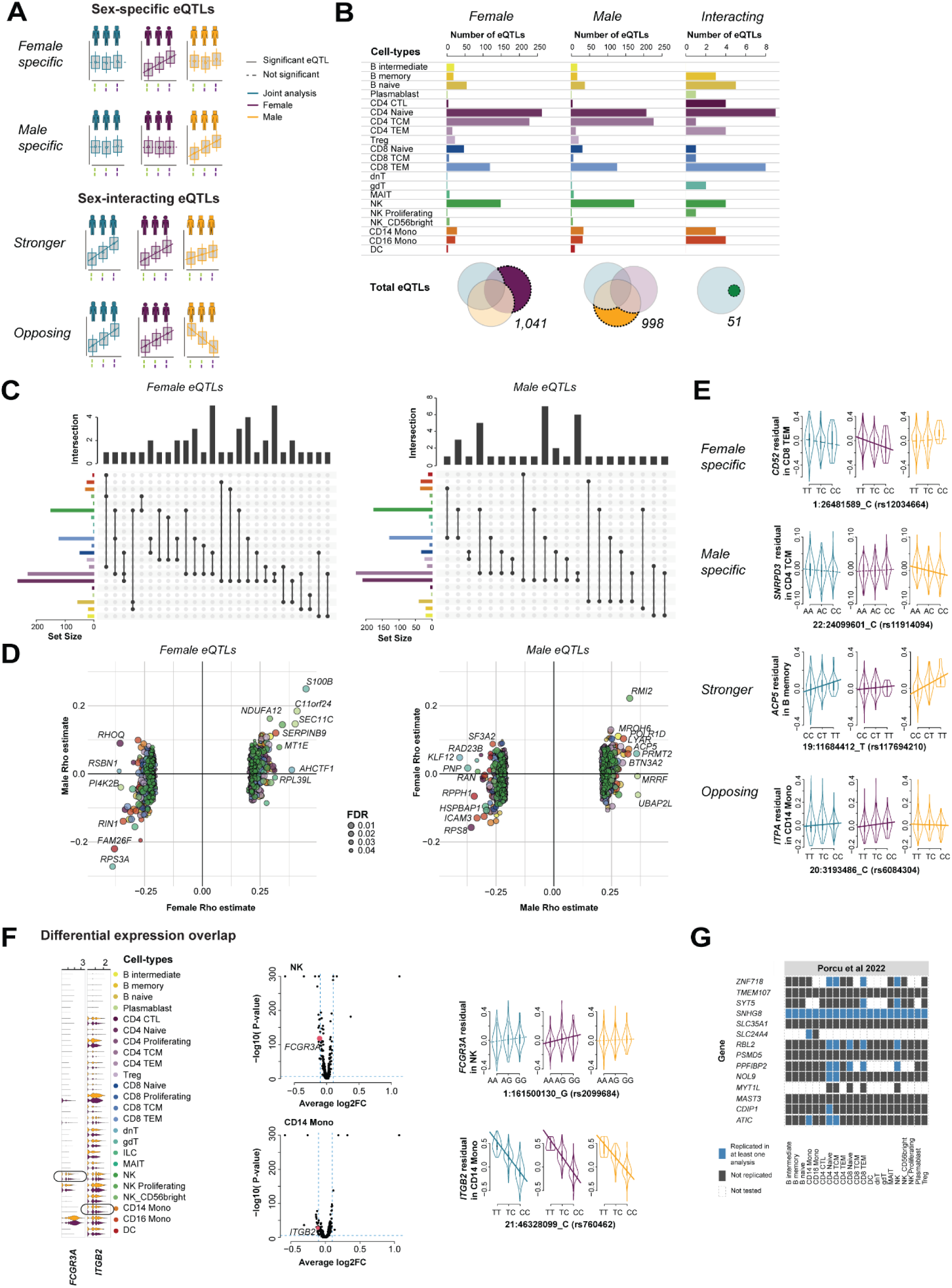
Sex-linked cis-eQTLs and sex-interacting eQTLs. (A) Sex-biased eQTLs can either be sex-specific or sex-interacting. Sex-specific eQTLs can be female or male, without significant association in the joint analysis or in the opposite sex. Sex-interacting eQTLs are significant in all analyses but have different effect sizes. (B) Total number of cis-eQTLs per cell type across the joint, stratified and interacting eQTLs. (C) Overlapping cis-eQTLs across cell types for female (left) and male (right) specific eQTLs. (D) Scatterplot of effect sizes (rho estimates) for female (left) and male (right) specific eQTLs plotted against estimate in the opposite sex. Colored by cell type and size of point reflects sex-specific FDR. (E) Example eQTLs for all four sex-biased eQTLs from panel A. (F) Overlap of sex-biased eQTLs with a sex-biased expression showing eQTL plot, differential expression, and expression of genes *FCGR3A* (female-specific) and *ITGB2* (interacting). (G) Replication of sex-interacting SNPs from Porcu et al. 2022 in our data per cell-type. Blue indicates replication, whilst gray boxes describe no replication. Dashed were not tested.

**Fig. 5:**
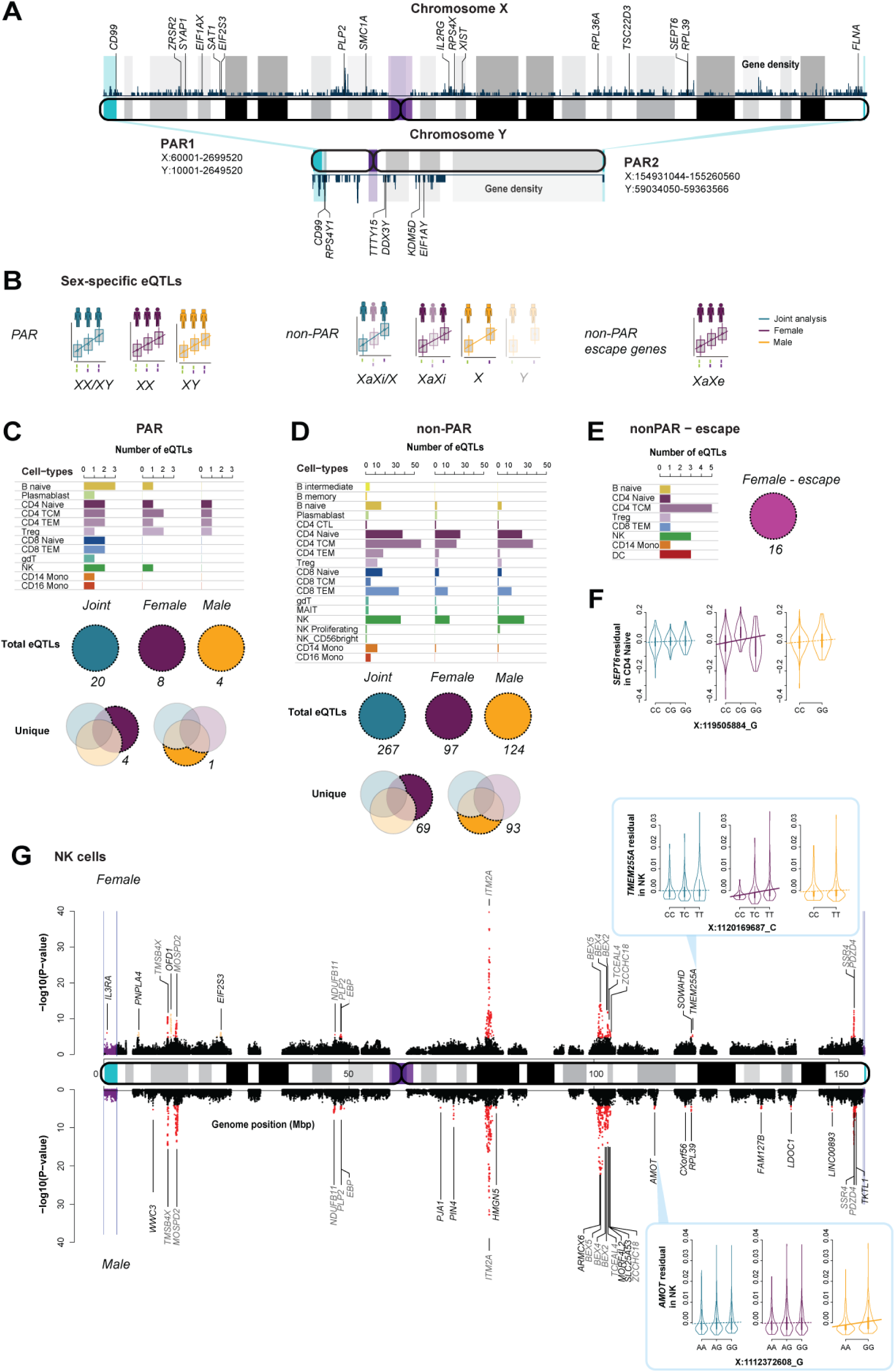
Sex chromosome eQTL analysis. (A) Ideograms of the X Y chromosomes showing PAR and non-PAR regions (cyan), and centromere (purple). Gene density (navy) histogram across the chromosomes, along with highlighted genes of interest on the X and Y. (B) Sex specific eQTL models of the sex chromosomes split into the PAR, non-PAR and X escapee genes. Each model highlights the joint analysis (turquoise), and the sex stratified (purple and gold). PAR analysis is similar to the autosomes, where genes escape X-inactivation. In contrast, heterozygous alleles in females in the non-PAR analysis require knowledge of the inactive/active allele (light-purple) as males are only ever homozygous. (C) Total number of cis-eQTLs by cell-type in the PAR regions, (D) non-PAR, and (E) non-PAR escape genes. (F) Example gene *SEPT6* with a female specific eQTL and female-biased DEG. (G) Example Manhattan plots of the NK cell. Top female, bottom males with female-specific (*TMEM255A*) and male-specific (*AMOT*) examples.

We next tested for functional enrichment of the eGenes with sex-specific associations. We found enrichment of transcription factor targets and immune-related functions from MSigDB(Liberzon et al., 2015a). Genes with associations in either females or males are likely to have sex-specific functions, not captured in the enrichment. For example, the female-specific eQTL of *CD52* rs12034664 in CD8 TEMs (**Fig. 4E**) may be linked to downstream activity: *CD52* plays a role as an effector molecule for suppression of Tregs in type I diabetes (Bandala-Sanchez et al., 2013). Interestingly, a second variant in DC (rs11577318) is a male-specific eQTL. These SNPs are not in LD (rs11577318 and rs12034664, R^2^=0.0087) and thus exhibit independent control of the same gene in different cell types. While *SNRPD3* rs11914094 in CD4 TCMs (**Fig. 4E**), a male-specific eQTL, is a subunit of the spliceosome, in mice this gene and associated complex have increased expression in male mice, believed to be linked to sex determination through alternative splicing (Planells et al., 2019).

### Sex-specific allele effects drive gene expression differences

The previous stratified analysis focused on cis-eQTLs that were absent in the other sex, yet there may be different effect sizes of the same variant which we call sex-interacting. To identify sex-interacting eSNPs (see **Box 1**), we took eQTLs from the joint analysis and tested for a genotype by sex interaction (see **Methods**). This analysis highlights expression differences that occur when the allelic effect of genotype differs between males and females (Kukurba et al., 2016). We identified 51 sex-interacting eQTLs (**Supplementary Table S13**). The majority (46 eSNPs) were in the same allelic direction, with 27 showing a stronger effect in males, and 19 in females.

#### Box 1 Definitions

*eQTL - expression trait quantitative loci*.

*Sex-specific eQTLs - cis-eQTLs that show female or male effects, but not both, when sex- stratified*.

*Sex-interacting eQTLs - cis-eQTLs that show opposing effects in males and females or weaker effects in one sex versus the other when sex-stratified. In some cases where we observe the eQTLs in the joint analysis but only in one sex, we have labelled these as ambiguous*.

*Autosomal eQTLs - sex-specific and sex-interacting cis-eQTLs that are tested on the autosomal chromosomes*.

*Sex chromosome eQTLs - sex-specific cis-eQTLs that are tested on the sex chromosomes. The analyses are split into PAR (diploid) and non-PAR (haploid) tests*.

*Sex-biased eQTLS - cis-eQTLs that show sex-specific effects or are sex-interacting from both the autosomes and sex chromosomes*.

Our approach was also able to detect effects present in just one sex or in the opposite direction of the allelic effect (5 eGenes). These include *NLRP2*, *ITPA*, *PQLC3*, *IL2RA* and *NAGK*. *NLRP2* (NACHT, LRR and PYD domains-containing protein 2, rs12969457 B naive) is a sensing component of NLRP2 inflammasomes and contributes to the regulation of immune responses regulating activities of NK-kB and acting as a pro-inflammatory molecule through caspase-1 activation(Tschopp et al., 2003). Furthermore, *NLRP2* also has reproductive functions, believed to be responsible for maintaining fertility in females (Kuchmiy et al., 2016) and establishing maternal–fetal tolerance during pregnancy (Tilburgs et al., 2017). *ITPA* (inosine triphosphate pyrophosphatase ITPase, rs6084304 CD14+ Monocytes) variants have been linked to chronic hepatitis C response treatment efficacy (Rembeck et al., 2014). *IL2RA* (interleukin 2 receptor subunit alpha, rs7261003, CD8 TEM) is involved in regulating immune tolerance by controlling regulatory T cells (Treg) activity (Chinen et al., 2016).

### Sex-biased gene expression in SLE-associated genes is influenced by genetic regulation

To link gene expression that is biased between sexes to genetic regulation, we looked for the overlap between the sex-biased eQTLs and the sex-biased differentially expressed genes. We identified most of the overlap signals occurring in eQTLs with joint effects (19 genes). However, we found one gene with a female-specific eQTL and sex-biased expression: *FCGR3A* in NK cells. This gene forms part of the immunoglobulin gamma (IgG) receptor and mediates IgG effector function in NK cells. It has also been associated with sex-linked traits: immunodeficiency of NK cells (Grier et al., 2012), including susceptibility to recurrent viral infections (Wang et al., 2017), severity of COVID-19 (Vietzen et al., 2022), and the autoimmune disease Systemic lupus erythematosus (SLE) (Zhu et al., 2016). More specifically, the rs2099684 variant is associated with Takayasu arteritis in the Han Chinese population, which has a 90% female bias(Chen et al., 2017). Other variants (rs396991) on *FCGR3A* are shown to affect the efficacy of antibody-dependent natural killer cell-mediated cytotoxicity in patients receiving rituximab treatment(Robinson et al., 2022a). Additionally, we find *ITGB2* with a sex-interacting eQTL in CD14+ Monocytes also showing female-biased expression in our data (**Fig. 4F**). *ITGB2* (Integrin beta chain-2), along with the alpha subunit, form integrin heterodimers involved in cell adhesion and cell-surface mediated signalling. It is linked to the inflammatory response in monocytes, along with roles in the autoimmune disease systemic sclerosis (scleroderma) (Xu et al., 2021). More recently *ITGB2* signalling pathway was shown to be enriched both in SLE and primary Sjogren’s syndrome (Cui et al., 2023). However, the specific variant (rs760462) has no known clinical correlation. Nevertheless, these two genes are also located in two distinct cis-regulatory elements and are interesting examples of sex-specific genetic regulation.

To verify and replicate our sex-specific eQTLs on autosomes, we looked for overlap with other bulk studies available (Kukurba et al., 2016; Oliva et al., 2020; Porcu et al., 2022; Yao et al., 2014). In these studies, little replication was evident between the studies themselves. Of the 26 genes identified overall, we were able to replicate 10 eGenes/eQTLs in a range of cell types with various analyses (**Fig. 4G**). Once again, this analysis suggests the ability of single-cell resolution to detect associations missed in bulk. Lastly, we conducted a gene co-expression network analysis, considering cell type and sex specificity, to identify additional genes sharing common regulatory control or pathway involvement (**Supplementary Fig. S3-5**). However, the generated networks did not overlap with modules for either *FCGR3A* or *ITGB2*.

### cis-eQTLs are depleted on the sex chromosomes

The effects of genetic variants on the sex chromosomes have not been assessed in many eQTL studies and may explain some of the sex differences. To assess the sex chromosomes for eQTLs, we separated our analysis of the X into PAR and non-PAR and then further into X escape and non-escape genes (**Fig. 5A)**. The non-PAR regions of the X chromosome are haploid, as females inactivate one of their X chromosomes, and males only have one copy of the X. This region spans most of the X chromosome, approximately 152Mbp. Around 953 genes are located in this region (GENCODE hg19/GRCh37), with 41 immune-related genes (∼4%) and approximately 66 genes escaping X-inactivation (Tukiainen et al., 2017). The pseudoautosomal regions (PAR1 and PAR2) are short regions of homology between the X and Y chromosomes at the tips of both chromosomes. The PARs are thus diploid, as genes on the female inactive X escape inactivation. In total, 26 protein-coding and lncRNAs sit in these PARs.

For the PAR analysis, we genotyped approximately 1,300 SNPs on the PAR XY, with around 300 passing QC. As with the autosomes, we ran our analyses both jointly and stratified (**Fig. 5B, Supplementary Table S16**). In the joint analysis, we found 14 eQTLs in PAR1 and 6 in PAR2, for a total of 20 eQTLs (9 eGenes). In the stratified analysis, we found eight eQTLs in females and 5 in males, all in PAR1. We identified no significant associations in PAR2, potentially due to loss of power. All eGenes detected in the stratified analysis were detected in the joint analysis (**Fig. 5C**) except *AKAP17A*, a protein kinase A anchoring protein, found in female Tregs (FDR∼0.0107)(Meester et al., 2020), and *IL3RA* in females in NK cells (FDR∼0.01). The latter encodes *CD123* (Interleukin 3 receptor alpha) and has been shown to influence COVID-19 responses between the sexes(Butler-Laporte et al., 2022).

In the non-PAR joint analysis, we found fewer eQTLs on the X chromosome relative to autosomes (**Fig. 5D, Supplementary Table S17**). We tested genes that do not escape X-inactivation by removing known X-escaping genes from our list (66 genes(Tukiainen et al., 2017), **Supplementary Table S14**). This was done to compare similar gene dosages across males and females. We found 97 significant eQTLs in the female stratified analysis. In males, we detected 124 eQTLs on the non-PAR X chromosome. The additional results in the male analysis were likely due to variation in XCI for hetereozygous females. We repeated the analysis jointly and found 267 eQTLs in total. Of the 97 eQTLs identified in females, 69 were unique (i.e., not in the joint and male analysis), and of the 124 in males, 93 were unique. And finally, we looked at genes that are known to escape X-inactivation in females. In this analysis, we found 16 eQTLs, with *XIST* and *RPS4X* being the most recurrent across cell types (**Fig. 5E**). Many sex-biased eQTLs were in similar eGenes, with differing lead SNPs. For example, in NK cells (**Fig. 5G**), 13 eGenes had all male-specific associations (*e.g.*, *AMOT*). At the same time, three were only female-specific (*e.g.*, *TMEM255A*) and 3 X-escape genes (*PNPLA4*, *OFD1* and *EIF2S3*) were not tested in males.

In addition to the X non-PAR, the Y chromosome has a non-PAR stretching 56Mbp, with 102 genes. In our data, we genotyped ∼7000 SNPs, with 3243 remaining following QC. As no recombination occurs on the Y, imputation here was difficult and unlikely to be informative. We attempted to use the haplogroups of the Y chromosomes(Poznik, 2016), however, all the males were classified as belonging to major haplogroups BT-M8947 and BT-M8949 that differ by one variant each and thus not enough variation to test for *cis*-eQTLs.

In the cases where genes are differentially expressed by sex on the sex chromosomes, this could likely be due to genetic control. To test this, we again looked for an overlap between the eQTLs and sex-DEGs. Of the PAR X chromosome genes, we observed no overlap of eGenes with sex-DEGs. In the non-PAR X, *SEPT6* was the only eGene with both female-biased expression and an eQTL in CD4+ Naive and CD4+ TCMs (**Fig. 5F**). *SEPT6*, a septin GTPase, plays a role in T cell migration(Dolat et al., 2014) and may potentially escape XCI(Shvetsova et al., 2019). Of the escape genes, *RPS4X, XIST,* and *EIF2S3* had both eQTLs and female-biased expression in a few of the cell types (B Naive, CD4+ Naive, CD4+ TCM, DC and Tregs). As in the autosomal analysis, no male-specific eQTLs overlapped male-biased expression. Together, these results indicate higher immune gene expression in females at baseline, reflected primarily in more female-biased expression but also alternate regulatory control, likely linked to X- inactivation and escape.

## Discussion

Our analysis of sex differences of PBMCs establishes the importance of studying the immune system in a sex-specific way. Despite known differences in the immune systems of males and females being recorded, many researchers still study and assess their functions, pathways, gene expression and genetic regulation agnostic of sex. This work examined these differences in this dataset of close to 1000 individuals and found that small cell-type proportion differences do exist between the sexes. These differences are likely to underpin functional effects - such as heightened responses to foreign stimuli (*e.g.*, monocytes) and autoimmune reactivity (*e.g.*, T and B cells).

In addition to differences in the cellular landscape of male and female individuals, we also identified sex-specific gene expression differences. Expectedly, many of these genes were on the sex chromosomes. However, the differentially expressed function of these genes is still open for investigation. Ribosomal genes that show male-biased expression could be linked to differences in proliferation, cell activation, and cellular exhaustion, which are common mechanisms in cancer cells. In contrast, the number of genes that showed sex-biased expression in females were linked to immune pathways, highlighting downstream activity likely to be influenced in activated or stimulated immune systems.

Through a comprehensive evaluation of the effect of sex and the X chromosome on control of gene expression in a cell type-specific manner, we identified numerous sex-specific eQTLs which were not previously observed in bulk whole-blood studies. The impact of genetic variation on gene expression on the X chromosome differed from autosomes, with smaller numbers of eQTLs overall and eQTLs with lower effect sizes. Moreover, the X chromosome eQTLs were less likely to be shared between cell types. These findings align with previous studies and support the hypothesis that a more efficient purification of selection on the X is present compared to autosomes(Kukurba et al., 2016). When considering the functional enrichment of sex-specific DEGs and eQTLs, we found that male-specific DEGs and eGenes were related to non-reproductive cancers. In contrast, female-specific ones were generally involved in immunological pathways.

However, our findings have their limitations. In addition to differences in sex, many other factors can play a role in gene expression. For example, immune responses change over a lifetime; as we age, B cells increase in females while regulatory T cells increase in males. Furthermore, there is considerable evidence that parity(Santucci-Pereira et al., 2019; Verlinden et al., 2005) and hormonal changes with menopause impact gene expression levels in mammary glands, which can change the risk of breast cancer. These changes may have also affected circulating immune cells and, therefore, should be considered in future studies. A further caveat to this analysis is that none of these cells were stimulated or activated by any infectious trigger, and the effect sizes we observe reflect baseline differences. In other conditions, these small differences may be exacerbated. Additionally, as we have taken our data at one time point, any fluctuations in hormone levels or dynamic/periodic changes to gene expression levels such as circadian rhythms, will also be missed or averaged away. Future work to assess the influence of infections, stress or other environmental factors may show additional or larger effect sizes.

There is a well observed disparity in disease prevalence between sexes with autoimmune diseases being more common in females(Klein and Flanagan, 2016a). Overall, our results suggest that genes with sex differences are involved in immunologically important functions, showing higher overall activity in females. These results highlight genes that are sexually dimorphic at baseline could potentially vary in their response in immune disease between the sexes in a cell-type specific way.

## Methods

### OneK1K data and cell-type classification

The OneK1K data set was generated at the GWCCG and described here(Yazar et al., 2022). Briefly, 1104 individuals from the Tasmanian Ophthalmic Biobank were genotyped and had PBMCs sequenced using the 10x genomics platform. Imputation was performed using the Michigan Imputation Server(Das et al., 2016), with Minimac4(Fuchsberger et al., 2015) and the Haplotype Reference Consortium (HRC) panel(McCarthy et al., 2016). Following quality control, 982 individuals remained, 564 females and 418 males. Sex was confirmed through a SNP-based sex analysis. Approximately 1200 cells on average were sequenced per individual, totalling 1,267,758 cells. A total of 75 batches (i.e., pools) were processed. Samples were multiplexed, between 10-14 samples per pool, and a total of 20,000 cells captured per pool. Sequencing was done with the Illumina NovaSeq 2000. Reads were processed using Cell Ranger Single Cell Software Suite (v 2.2.0; 10x Genomics)(Zheng et al., 2017), followed by demultiplexing into their respective pools. Mapping and alignment were done to GRCh37/hg19 (release 84) using STAR(Dobin et al., 2013) within the Cell Ranger Suite. Batch correction and SCT transformation were performed through Seurat(Hao et al., 2021a). Cell-types were classified using the Azimuth pipeline with the human PBMC reference for L1 and L2(Bakken et al., 2021). Dendritic cells classed as ASDC, cDC1, cDC2 and pDC were merged due to low cell numbers. Other rarer cell-types were retained but were removed from specific analyses when more cells were required.

### Cell-type proportions

For each individual, cell-type proportions were calculated based on total-cell count / cell-type. These were then adjusted using the arcsin transformation from the Speckle R package(Phipson et al., 2022). The *propeller* function was used to determine if the average proportions were significantly different between the sexes using an F-test.

### Age regression analysis

We tested for significant correlation between cell proportions and age using Spearman’s rho and the adjusted p-value (*cor.test* in R). This was done per cell-type, first jointly across the sexes, and then stratified by sex.

### Sex-differential expression analysis

We ran single-cell differential expression analysis for each cell-type with the *FindMarkers* function in Seurat using the Wilcoxon test, with default parameters and log2FC threshold set to 0. We then filtered on significance using an adjusted P-value (FDR) of 0.05 and |log2FC| greater than 0.1.

### Sex chromosome gene expression analysis

Using sex chromosome genes on the X and Y chromosomes as the set of variable genes, we ran dimension reduction (UMAP) through the Seurat R package(Hao et al., 2021a) to delineate cells based on sex. We visualized these results via their UMAP dimensions with the *DimPlot* function in Seurat. We highlighted the expression of a subset of genes of interest using the *plot_density* function in Nebulosa(Alquicira-Hernandez and Powell, 2021).

### Cell-type gene marker identification

For each cell-type, we determined cell-type marker genes using the *FindAllMarkers* function in Seurat with a |log2FC| greater than 0.25 and percent expressed 0.25. We then filtered on significance using an adjusted P-value (FDR) of 0.05. We performed this per batch and took the genes that recurred in at least 80% of the 75 batches as markers. We repeated this in a sex-stratified manner to obtain cell-type markers that were conditioned on sex.

### Classifier and class prediction

To test for the ability of the gene sets to label or classify cells, we use a “classification” score based on the ranked gene expression levels and calculated an enrichment statistic (*analytic_auroc* in EGAD (Ballouz et al., 2017)). We expect high scores (close to 1) for gene sets that are highly expressed within cells (i.e., markers).

### Gene sets and functional enrichment analysis

We curated gene sets from multiple biological domains. We downloaded the Gene Ontology (GO)(Ashburner et al., 2000; Gene Ontology, 2021) and the generic GO slim subset. Additionally, we used MSigDB (Subramanian et al., 2005), with a focus on the HALLMARK (Liberzon et al., 2015b), KEGG (Kanehisa and Goto, 2000), REACTOME(Gillespie et al., 2021) and BIOCARTA (Nishimura, 2001) gene sets and pathways. We downloaded regulatory TFs from MotifMap (Rouillard et al., 2016) and ENCODE TFs-target (Consortium, 2011) gene sets curated through Harmonizome(Rouillard et al., 2016). Furthermore, we generated curated X-linked datasets from sex-differential and X-inactivation analysis papers. We labelled these datasets as: Jansen2014 (Jansen et al., 2014), Mele2015 (Melé et al., 2015), Tukiainen2017 (Tukiainen et al., 2017), Schmiedel2018 (Schmiedel et al., 2018), Bongen2019 (Bongen et al., 2019), and Oliva2020 (Oliva et al., 2020).

For gene set enrichment analysis, we used the hypergeometric test in R (*phyper*) and adjusted for multiple tests using *p.adjust*. For network assessment and analysis, we ran the neighbour-voting algorithm in EGAD in R(Ballouz et al., 2017) which uses the guilt-by-association (GBA) principle to assess network connectivity.

### Sex-specific eQTL and sex-interacting eQTL analysis

We ran cis-eQTL analysis per cell-type across all the autosomes jointly (code available from https://github.com/powellgenomicslab/onek1k_phase1). Details of this analysis were described previously(Yazar et al., 2022). In brief, average expression of each gene per person across all genes available for each cell type was calculated using the corrected counts with SCTransform(Choudhary and Satija, 2022). We then calculated the number of individuals with non-zero expression for each gene and filtered genes expressed in less than 10% of the cohort. All values are then log transformed (log x+1). Within each cell type, cis-eQTLs were identified by Spearman’s rank correlation testing using residual expression levels adjusted for sex, age, first four genotype-based principal components (PCs) and two PEER factors from original analysis. We restricted our search to variants within 1Mb of the TSS of either end of a gene. The resulting SNP-gene pairs were filtered at the FDR threshold of 5% at the chromosomal level for each cell type and the most significantly associated SNPs were labelled as cis-eQTLs.

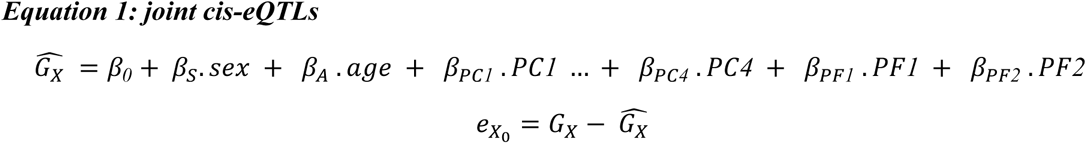

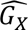 is a matrix consisting of the average expression of gene X per individual 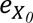. is the matrix including residual expression of gene X after adjusting for sex, age, six genotyping principal components, and two PEER factors.

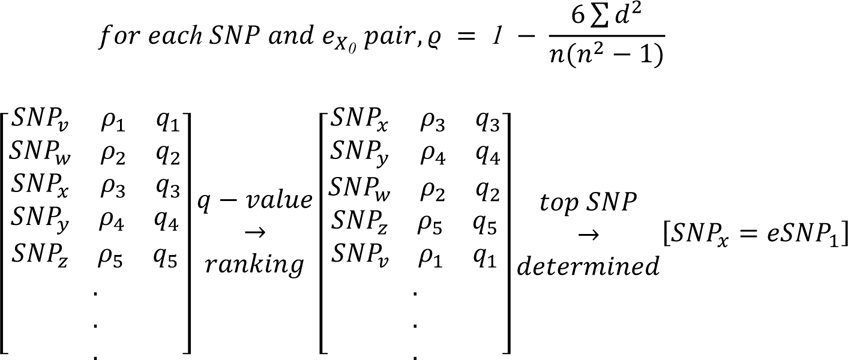

𝜌 is the correlation between 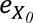 and a matrix of three genotypes coded as 0,1 and 2 where 2 represents the assessed allele for each of 5,433,038 SNPs and *q* is the associated *q*-value. *d* is the difference between two rankings of residuals (*e_A_*) and *n* is the number of measurements.

Then using these cis-eQTLs results, we ran a sex-interaction analysis to identify eQTLs with varying effect sizes by sex. In this analysis, for each gene-SNP pair within each cell type, we fit a linear regression model and tested for genotype-by-sex interaction while adjusting for previously mentioned additional factors:

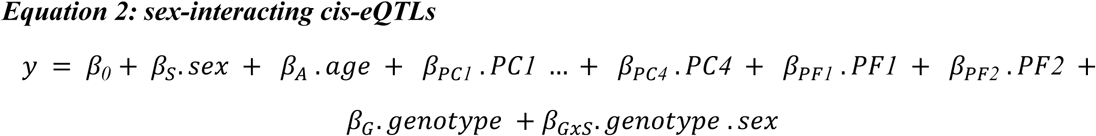

where *y* is the gene expression, 𝛽*_0_* is the intercept and 𝛽 are the corresponding effect sizes. 𝛽_𝐺𝑥𝑆_ is the effect size of genotype-by-sex interaction on gene expression. Since we have already applied a multiple testing correction and accounted for the number of independent eQTLs tested per chromosome in our initial analysis, we applied Storey q-value across genes to identify genes with at least one significant (FDR ≤ 0.25) sex-interacting eQTL.

We also repeated our original analysis in a sex-stratified manner, using age, first four genotype-based principal components (PCs) and two PEER factors for each sex.

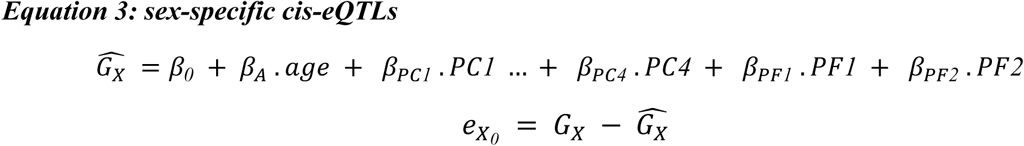

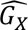 is a matrix consisting of the average expression of gene X per individual. 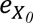 is the matrix including residual expression of gene X after adjusting for age, six genotyping principal components, and two PEER factors.

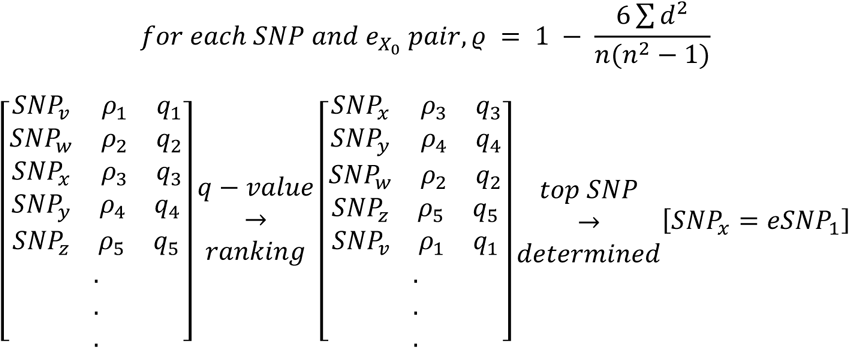

𝜌 is the correlation between 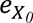 and a matrix of three genotypes coded as 0,1 and 2 where 2 represents the assessed allele for each of 5,433,038 SNPs and *q* is the associated *q*-value. *d* is the difference between two rankings of residuals (*e_A_*) and *n* is the number of measurements.

In the OneK1K cohort, we have slightly more female participants than males. To ensure equal power to detect eQTLs in each sex, we downsampled the number of female participants in each cell type and matched to the male participant numbers. After applying FDR threshold of 5% at the chromosomal level for each cell type and identifying the most significant cis-eQTL per gene per cell-type in each sex, we implemented two additional statistical assessments to filter away false positives. In our first level of testing, we first subsetted cis-eQTLs identified in the female analysis only, then assessed the distribution of corresponding p-values of the same SNP-gene pairs from the male analysis results and estimated the proportion of true null p-values (pi0) (Storey and Tibshirani, 2003). Then we applied this threshold to remove SNP-gene pairs from the list of female eQTLs. In our second level of testing, we used the two-sample z-test to compare the beta estimates of two populations to see if it is feasible that they come from the same population. For this analysis, we ran the stratified analysis using *MatrixEQTL*(Shabalin, 2012) and generated beta estimates and standard errors for each SNP-gene pair in each cell type for females and males. Next, we calculated the z-statistics for cis-eQTLs identified in each sex analysis:

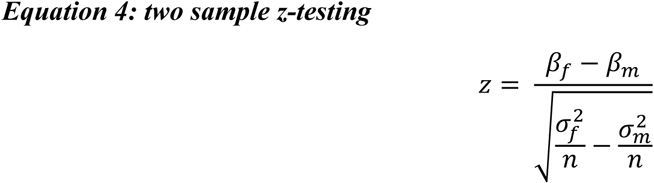

and we filtered out cis-eQTLs at the p-value of 0.05 threshold. In our final step, we defined a sex-specific eQTL only if the cis-eQTL passed both levels of testing.

### Imputation of the sex chromosomes

The sex chromosomes in humans share pseudoautosomal regions (PAR) and are assessed as diploid regions. PAR1 (X 60,001 to 2,699,520 and Y 10,001 to 2,649,520) and PAR2 (X 154,931,044 to 155,260,560 and Y 59,034,050 to 59,363,566) are located at the tips of the chromosomes and recombine. The remainder of each chromosome is labelled the non-pseudoautosomal region (non-PAR). In females, the non-PAR X is genotypically diploid, while in males the non-PAR Y is haploid. Thus, to analyse the sex chromosomes, we needed to impute them split by these regions. For the X chromosome, we extracted genotyped SNPs from the genotype PLINK file from the non-PAR (“chrX or 23”) and PAR (1 and 2) (“chrXY or 25”). Further filtering was performed on the PAR variants to match the Haplotype Reference Consortium (HRC) panel. Imputation was performed using the Michigan Imputation Server (Das et al., 2016) with Minimac4 (Fuchsberger et al., 2015) and the HRC panel (McCarthy et al., 2016), separately for the non-PAR and PAR segments. For the Y chromosome, we extracted genotyped SNPs from the genotype PLINK file (“chrY or 24”). As the non-PAR Y does not undergo recombination like the non-PAR X does in females, we cannot impute genotypes on the Y. Instead, we can haplotype the Y based on their genotypes. To this end, we ran *yhalpo* (Poznik, 2016) to identify the broad haplogroup of the male individuals.

### Sex-chromosome eQTL analysis

We performed first sex-stratified then joint eQTL analysis for PAR1, PAR2 and non-PAR regions separately. When PAR1 regions were tested, both genotypes on chromosome X and Y were modelled as 0 (homozygous for allele 1), 1 (heterozygous for allele 1) and 2 (homozygous for allele 2) and the analysis was performed as described for sex-specific eQTLs (**Equation 3**). Due to differences in gene locations on X and Y chromosomes, we tested an average of 4419 SNP-gene pairs per cell type in females and 4318 SNP-gene pairs per cell type in males. Joint sex-chromosome eQTLs were identified as depicted in Equation 1 where sex was included as a covariate when calculating the residuals before testing for Spearman’s rank correlation. To overcome the differences in available SNP-gene pairs between females and males, only pairs that were present in both sexes were included in this analysis. We conducted the eQTL analysis for PAR2 region in the same manner as PAR1.

Similar approach was followed when analyses were performed for non-PAR region with the difference being when testing for males, genotypes on chromosome X were modelled as 0 (homozygous for allele 1) and 1 (homozygous for allele 2) and analysis was completed using non-parametric Mann-Whitney U-test.

### Co-expression network generation

For each batch, we generated individual expression matrices using the normalized count data for each cell-type and sex. We used the most recurrent genes across all the data to ensure a common gene reference, totalling 14,716 genes. Then, we built a gene co-expression network from each of these expression matrices, using Spearman’s correlation to measure gene-gene relationships. Due to low cell-numbers for some cell-types, we built networks for 21 of the 27 cell-types. Each network was then ranked to standardize the correlations. We then aggregated the same cell-type per sex to build meta-analytic co-expression networks. We also built individual cell-type specific networks that were not conditioned on sex, and similarly aggregated these networks (labelled “joint”). For additional analyses, we then built “PBMC’’ aggregates from the cell-type aggregates, for the joint and individual sexes. Finally, we built networks from each individual, removing Erythrocytes, HSPC and Platelets. We aggregated individuals by their genotyped sex. Overall, we generated 70 aggregate networks.

### Differential co-expression analysis

To compare across networks, we assessed network differences based on overall topology, node degrees and functional performances. For each network, we took the correlation of the matrices as a measure of topological similarity. We calculated the node degrees of each network by summing the weights of each node, and then took the correlation of these node degrees. And finally, using the outputs from the GO slim analysis, we calculated the similarity of their performances by taking the correlation of the AUROCs of the functional groups. To identify the changes by genes and gene modules, we calculated the frequency of genes whose co-expression ranking changed by at least 0.5 between the networks. With these tallies, we identified which genes had the most topological shifts. Secondly, we repeated this with the node degree of the genes, calculating standardised residuals and selecting outliers that were three standard deviations away.

## Acknowledgements

This research was supported by a National Health and Medical Research Council Research Fellowship (S.Y.), Leader Fellowship (A.W.H., 2009079), Career Development Fellowship (J.E.P., 1107599), and Investigator Fellowship (J.E.P., 1175781). K.A.F. is supported by the Alex Gadomski Fellowship, funded by Maddie Riewoldt’s Vision. Additional grant support was provided by the National Health and Medical Research Council (1150144,1143163 and 2020517), the Australian Research Council (180101405), and the Royal Hobart Hospital Research Foundation.

The content is solely the responsibility of the authors and does not necessarily represent the official views of the funding agents. The funders had no role in study design, data collection and analysis, decision to publish, or preparation of the manuscript.

## Data availability

OneK1K single-cell gene expression and genotype data are available via Gene Expression Omnibus (GSE196830 https://www.ncbi.nlm.nih.gov/geo/query/acc.cgi?acc=GSE196830). The cell by gene data are available at Human Cell Atlas (HCA) (https://cellxgene.cziscience.com/collections/dde06e0f-ab3b-46be-96a2-a8082383c4a1).

## Code availability

The analysis code can be accessed at the GitHub repository (https://github.com/sarbal/sex_diffs).

## Supplementary Tables

**Supplementary Table S1. Cell-type counts and proportions**

**Supplementary Table S2. Correlation of cell-type proportions with age**

**Supplementary Table S3. Sex DEGs summary results per cell-type**

**Supplementary Table S4. List of sex DEGs per cell-type**

**Supplementary Table S5. List of gene markers per cell-type**

**Supplementary Table S6. Gene markers per chromosome enrichment analysis**

**Supplementary Table S7. Gene markers overlaps with sex DEGs**

**Supplementary Table S8. Aggregate co-expression network clustering analysis**

**Supplementary Table S9. Sex- specific eQTL summary results – autosomal**

**Supplementary Table S10. Sex- specific eQTL results – autosomal – joint**

**Supplementary Table S11. Sex- specific eQTL results – autosomal – female**

**Supplementary Table S12. Sex- specific eQTL results – autosomal – male**

**Supplementary Table S13. Sex- specific eQTL results – autosomal – interacting**

**Supplementary Table S14. List of X-escape genes**

**Supplementary Table S15. List of par genes**

**Supplementary Table S16. List of MHC genes**

**Supplementary Table S17. Sex- specific eQTL summary results – sex chromosomes**

**Supplementary Table S18. Sex- specific eQTL results – PAR**

**Supplementary Table S19. Sex- specific eQTL results – non-PAR**

**Supplementary Table S20. Sex- specific eQTL and sex DEGs overlaps counts**

**Supplementary Table S21. Sex- specific eQTL and sex DEGs overlap gene list**

## Supplementary Figures

**Supplementary Fig. S1 Cell-type proportions and age changes**

**Supplementary Fig. S2 Sex-DEG enrichments in other gene sets**

**Supplementary Fig. S3 Co-expression by cell-type and sex**

**Supplementary Fig. S4 Co-expression downsampling analysis**

**Supplementary Fig. S5 Differential co-expression comparisons**

**Supplementary Fig. S6 Functional enrichment results of aggregate networks using EGAD across gene sets**

