## Supplementary text for "The impact of sex on the immune system explored at the single-cell level"

***Impact of age on sex differences in immune cell types***

Cell type abundances change with age, and this occurs in a sex-specific manner(Klein and Flanagan, 2016; Márquez et al., 2020). The sex-specific changes in cell proportions with age is linked to hormonal changes during puberty, menstruation and menopause(Taneja, 2018) and loss of clonal diversity in stem cell populations(Mitchell et al., 2022). When comparing proportions to recorded biological age **(Supplementary Fig. S1, Supplementary Table S2**), we see clear correlation patterns in NK cells (Rho=0.32, FDR~0.002)(Hazeldine and Lord, 2013) and CD8+ Naïve T-cells (Rho=-0.67, FDR< 0.001)(Chen et al., 2013), both of which change with age irrespective of sex. However, we see no significant correlation with Tregs with age which had been reported(Churov et al., 2020), instead NK proliferating cells, Plasmablasts, CD8+ TEM and gamma-delta T cells (gdT) are correlated with age in females (Rho NK proliferating=0.17 FDR~ 0.001, Rho CD8+ TEM=0.23, FDR~3.7x10^-7^ and Rho gdT=-0.25, FDR~4.4x10^-8^) but not in males.

#### ***Cell-type and sex-specific co-expression replicates known biological functions***

In addition to genetic control, we also wished to check whether observed sex-biased expression was linked to co-variation (*i.e.,* co-expression or co-regulation). Genes that co-vary are believed to be co-functional, either under common regulatory control or are in common pathways. Like gene set enrichment analyses, this approach includes additional network information in the enrichment, defined as a gene-gene interaction. Thus, analysing co-expression gene-gene networks that are conditioned on properties of the underlying cell types, tissues or sex highlights modules or interactions that are specific to these conditions. To assess sex-specific networks derived from PBMCs, we built cell-type and sex-specific networks from the data. We performed differential co-expression to determine the sex and cell-type specific modules and gene sets that differ by sex and/or cell type (**Supplementary Fig. S3**). To perform this robustly and remove confounding variables such as batch effects, we built separate networks per pool and aggregated these networks(Ballouz et al., 2015). By conditioning on sex and cell-type, we aimed to remove co-variation confounded with these conditions and capture the intrinsic variation within each “state”. We use the performance of the neighbour-voting algorithm in cross-validation (EGAD(Ballouz et al., 2017), **Methods**) to measure enrichment for known biological pathways and gene sets, with model performance measured by the averaged AUROC (area under the ROC curve) for each group across the *n*-folds (**Supplementary Fig. S3A**).

Using the Gene Ontology (GO (Ashburner et al., 2000; Gene Ontology, 2021)) slim gene sets to measure broad biological functions first, we find similar average AUROC scores between the sexes of the same cell-type, with average scores varying based on cell-type (**Supplementary Fig. S3B,** AUROC~0.53-0.58). The joint aggregate networks show slightly higher performances than the sex-stratified networks, even with down-sampling, suggesting that some of the connections within these joint networks are lost when conditioning on sex – *i.e.*, sex has some influence on co-expression and contributes to the broad GO pathways and functions (AUROCs +0.02). We see that the individuals’ network aggregates outperform the cell-type specific networks (AUROC~0.6), highlighting that cell-type composition (variation) drives a fraction (+0.04) of the additional co-expression in these networks. Notably, these performances are still just above average, reflecting the sparsity in single-cell RNA-seq data, as many genes are not expressed or detected and could not be assessed. The co-expression analysis reaffirms, as in bulk data, that a large fraction of the co-expression we observe is driven by cellular composition (Farahbod and Pavlidis, 2020). Conditioning on cell type removes these connections in the network, impacting the AUROC performance. Sex, on the other hand, has much less of an effect, as removing sex-specific connections in the network does not impact the AUROC as much.

We then looked at the specific gene set module differences between the networks (**Supplementary Fig. S3C**). The highest performing gene set module in most of the cell-types was GO:0003735 (structural constituent of the ribosome, AUROC~0.79), yet had lower performances in NK proliferating and plasmablast cells (AUROC~0.61). On the other hand, NK proliferating and plasmablast cells co-express mitotic-related genes (GO GO:0000278, mitotic cell cycle AUROCs~0.75), while other cell types do not (AUROCs~0.5). To identify functional specificity further, we ran our analysis on additional gene sets from the Molecular Signature Database (MSigDB(Subramanian et al., 2005)), with a focus on the HALLMARK(Liberzon et al., 2015), KEGG(Kanehisa and Goto, 2000), REACTOME(Gillespie et al., 2021) and BIOCARTA(Nishimura, 2001) gene sets and pathways. As before, we see cell-type specific networks scoring highly in particular pathways and gene sets related to their cell-type specific functions. Of note were monocyte-specific networks, where higher AUROCs in pathways linked to hypoxia, apoptosis and TNFɑ signaling were similar to the pathways found to be sex-differentially expressed in those cell-types (**Supplementary Fig. S6**). This suggests that these pathways and their genes show differential activity in monocytes (So et al., 2021; Varghese et al., 2022) dependent on sex.

In addition to pathways, co-expression may reflect co-regulation. Thus, we tested for TF-target enrichment within the networks, and whether we could observe differences in co-regulation between the sexes. We used the Gene Transcription Regulation Database (GTRD)(Kolmykov et al., 2021). TF-target genes dataset curated in MSigDB and observed highly correlated performances between the sexes once more (**Supplementary Fig. S3D**). The biggest differences were in the co-expression values of the target genes of *NR5A1* (steroidogenic factor 1 (SF-1)), a transcriptional activator involved in sex determination, and *CREBL2*(Ma et al., 2011), linked to adipose tissue differentiation. Whether these genes play roles in the immune system requires additional validation, but their roles in sex differences and sex-specific phenotypes (such as adipogenesis) are well established.

To further explore the performance differences, we looked at the topological network differences (i.e., gene-gene connections, **Supplementary Fig. S3E**) by comparing the pairwise co-expression ranked values of the cell-type specific aggregate networks. On average, ~16% of the edges change between the networks, with the most differences between the CD4+ T cell aggregates and the other cell types (~38%). Overall, we see that sex differences on average are low, as most cell-type specific networks have greater similarities to their same cell-type or similar cell-type within the hematopoietic hierarchy. To further measure the impact of sex and cell-type conditioning on the network topology, we then compared the change in overall connectivity per gene by assessing the node degree changes. As in the previous analysis, node degrees were similar between cell-type specific aggregates conditioned on sex, with most differences appearing between the different cell types (**Supplementary Fig. S3F**). On average, less than 1% of genes (~99) have changes in their node degree standardized residuals of 3SDs across cell types (holding sex constant). In contrast, between the sexes of the same cell type, there are less than ~0.5% of genes that have significant node degree differences (~43 genes). We find these genes to be enriched for those on the sex chromosome, a few of which are paralogs (e.g., *UTY* and its X paralog *KDM6A/UTX*, *ZFY* and *ZFX,* *KDM5D* and [*KDM5C*](https://www.genecards.org/cgi-bin/carddisp.pl?gene=KDM5C)). X chromosome paralogs are likely more variable as they are known to escape X-inactivation. Overall, we observe little sex network differences from the autosomal genes, reaffirming our observation of few detectable changes in these networks as reflected by their similar AUROCs performance. This analysis highlights that sex differences have small effects on the broad biological pathways or functions in contrast to cell type, as highlighted by the more similar AUROCs in the former and specific gene set performances differences in the latter.

We identified core co-expressed gene-gene modules in the immune cell types along with gene expression differences. Our work highlighted differential co-expression changes between the sexes primarily a result of sex-chromosome genes, along with a few autosomal genes. The regulatory relationships that were impacted were not significant, but the differences do reflect changes to signalling pathways that are likely buffered in a sex-specific way. The Y chromosome is known to influence inflammatory pathways. As such, females have paralogs to several Y-specific genes that escape X-inactivation to compensate. Their co-expression partners and regulatory factors are likely shared, which we observe in our data. RNA-seq analysis may need to distinguish between the XY paralogs and hence expression estimates, and subsequently, co-expression relationships may be obscured or miscalculated.

### **Supplementary Figures**

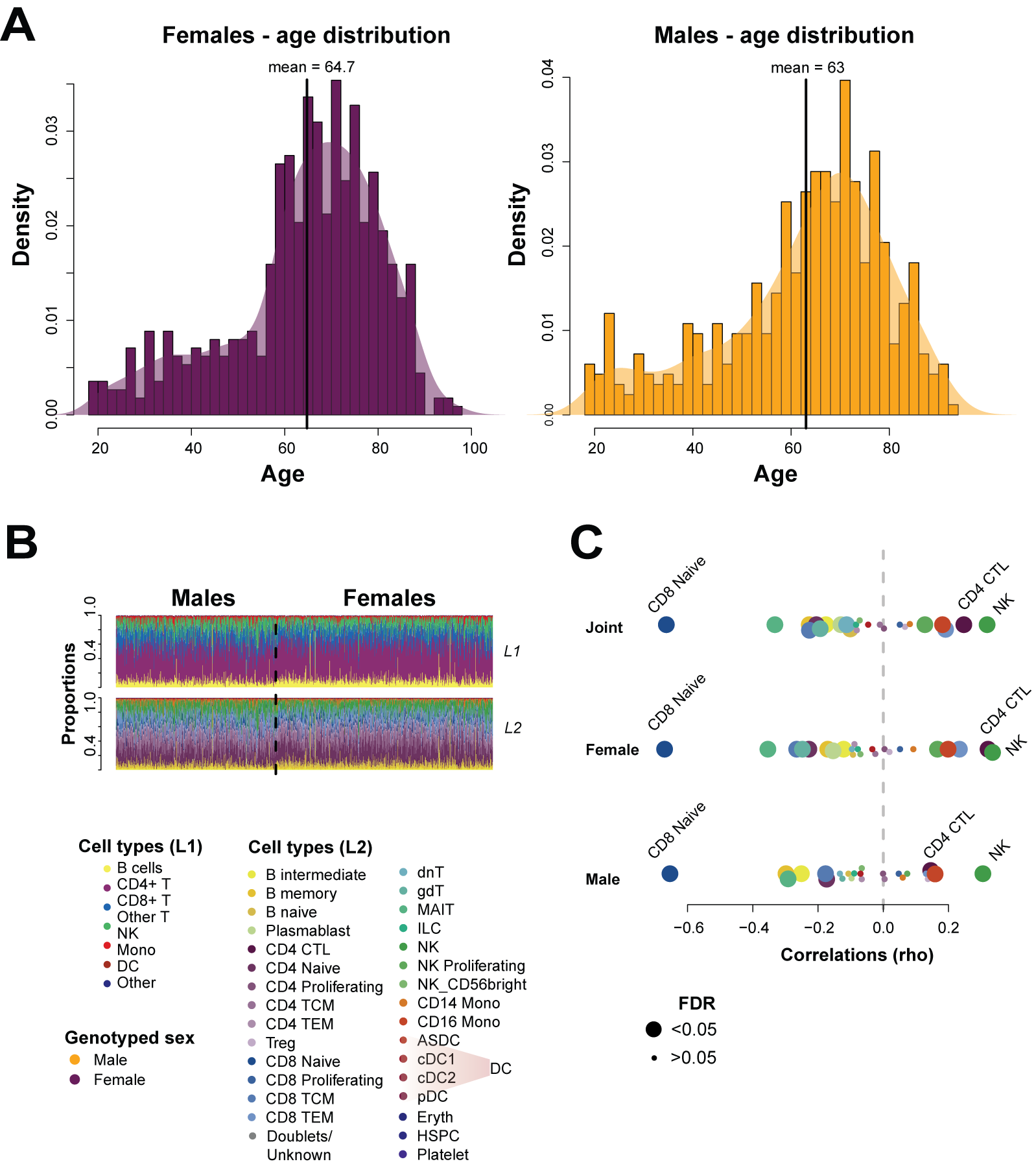

**Supplementary Fig S1 Cell-type proportions and age changes.** (A) Distribution of ages for females and males in OneK1K. (B) Distributions of cell-type proportions for L1 (broad) classification and L2 classifications across all individuals. (C) Correlations of cell-type proportions to age for both sexes (joint), females only and males only.

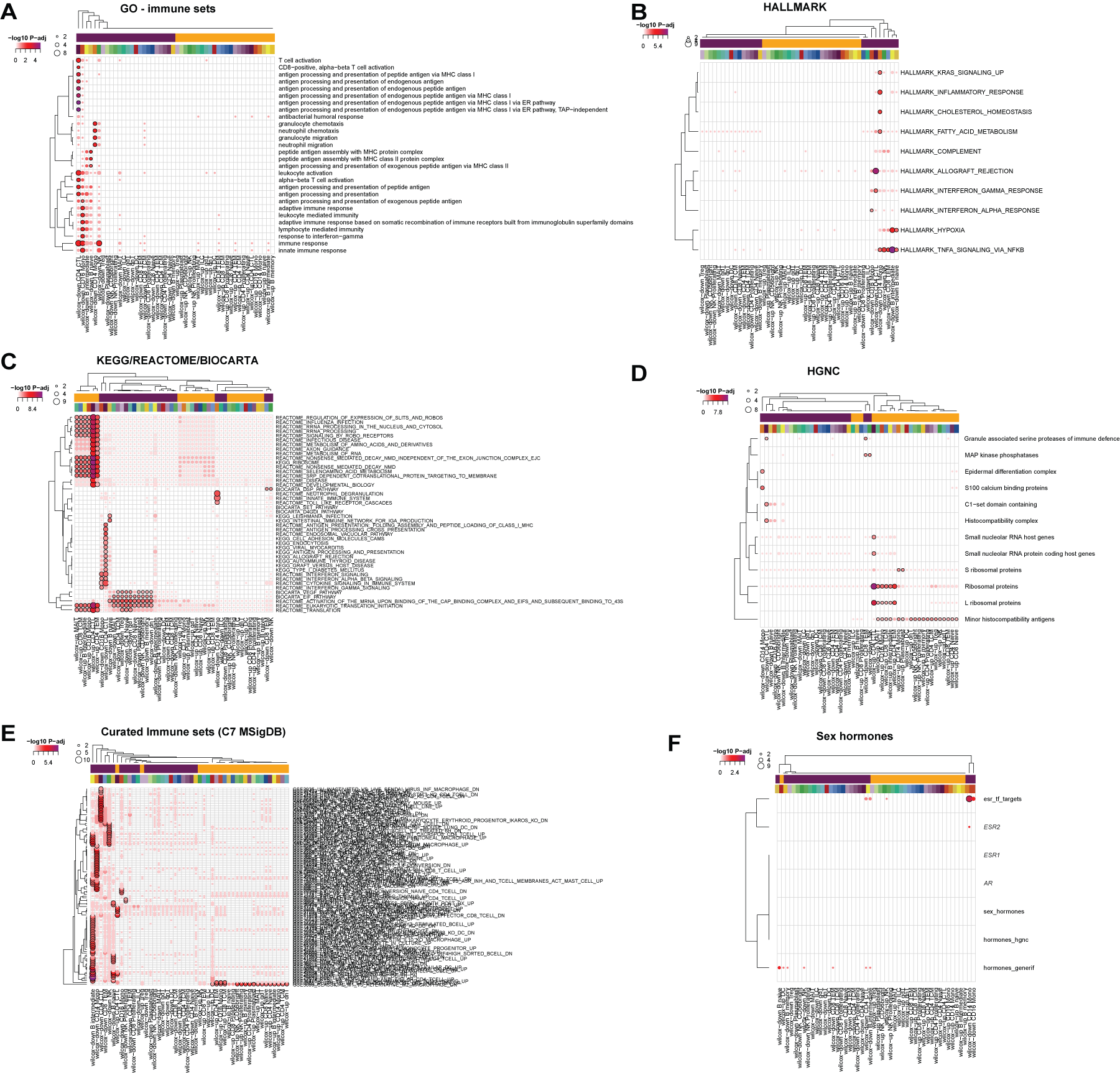

**Supplementary Fig S2 Sex-DEG enrichments.** Gene set enrichment analysis for (A) GO immune sets, (B) HALLMARK, (C) KEGG/REACTOME/BIOCARTA, (D) HGNC, (E) Curated immune gene sets (MolSigDB C7) and (F) sex hormones and their targets.

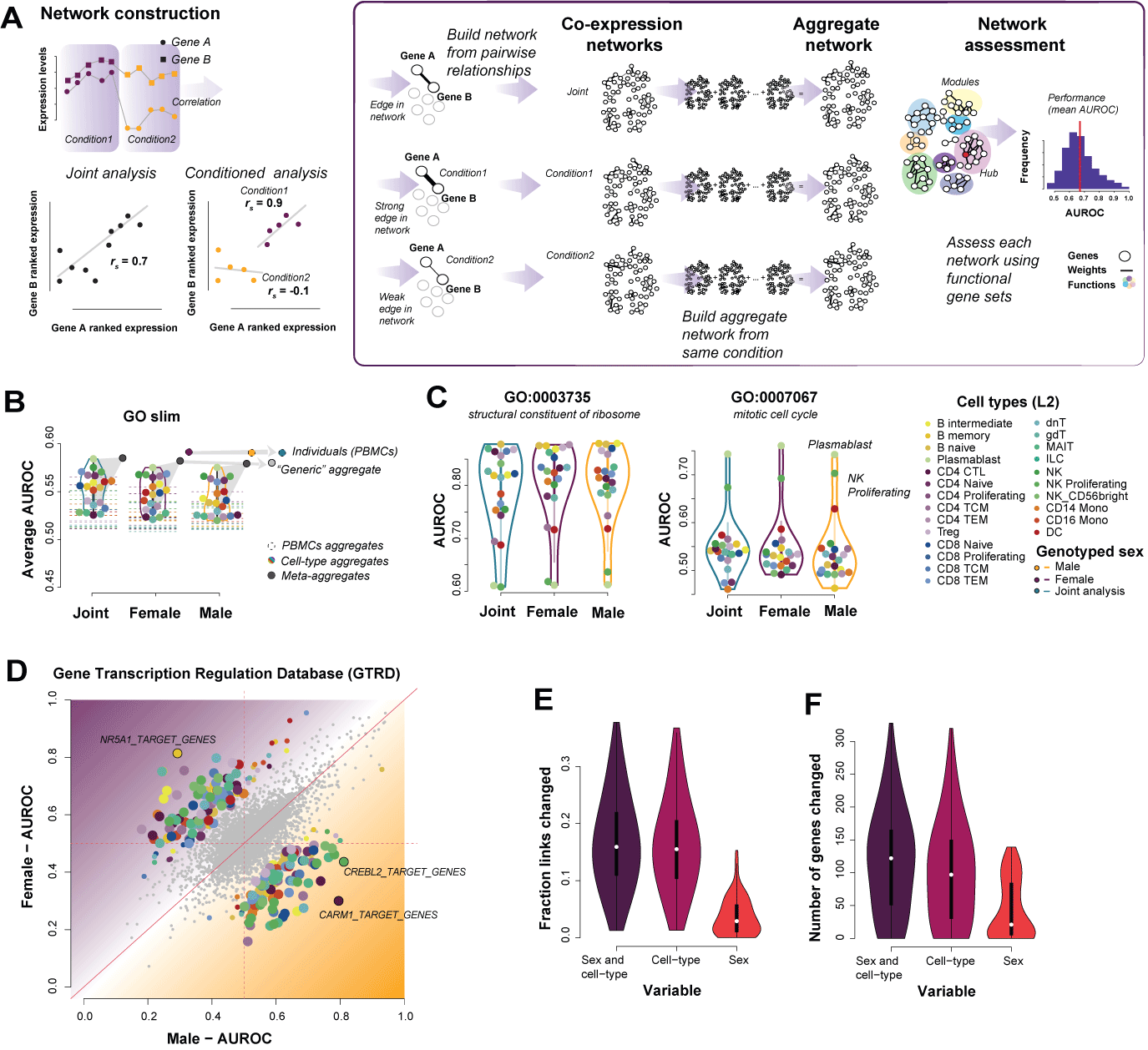

**Supplementary Fig S3 Co-expression by cell-type and sex.** (A) Building co-expression networks and assessing performance. For each subset of cells, we calculate pairwise correlations between genes, which generates an edge in the network. This is repeated for different conditions (sex and cell-type), and these individual networks are aggregated into a final cell-type specific and sex-cell-type specific network. Enrichment of pathways and gene functions within modules formed in the network are tested through our neighbour-voting algorithm and recorded as an AUROC performance. (B) Network aggregation and performances of cell-type and sex-specific aggregates with GO slim. (C) Gene set specific performances showing differences for cell-types in ribosome and mitotic specific pathways (D) Comparing male AUROCs and female AUROCs for TF-target gene performances. Comparison of aggregate networks based on ranked expression weights (E) and node degree (F). On average, when comparing within the same cell-type but varying sex, we observe lower impacts on the edges, while comparing across cell-types within the same sex or across sex, the average number of differences are higher.

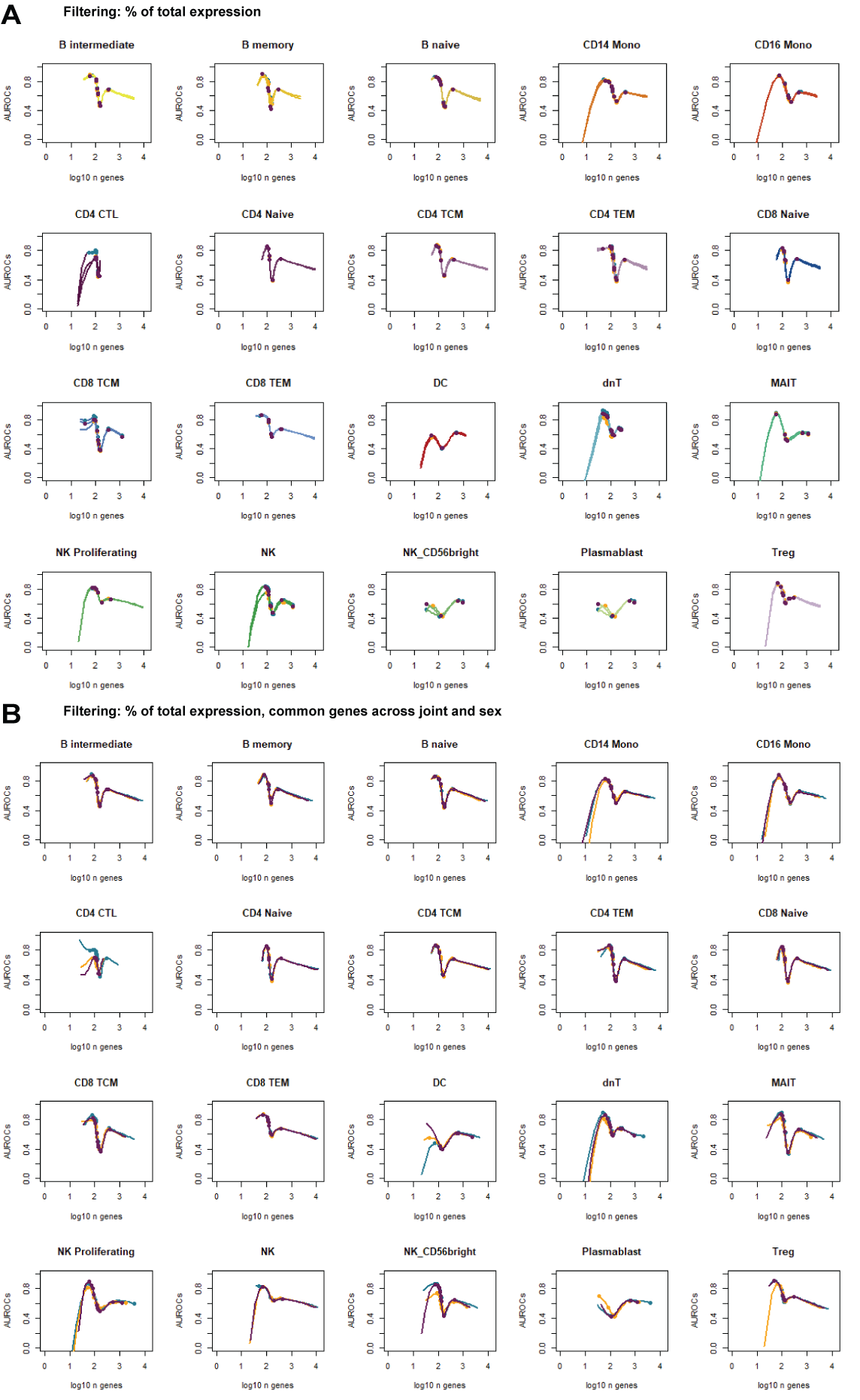

**Supplementary Fig S4 Aggregate co-expression downsampling comparisons.** (A) For each cell-type aggregate, we filtered genes that were expressed in a fraction of the cells in that specific parameter set, (B) and those that overlapped across the sexes or joint analysis. The x-axis shows the size of the final set of genes (log10), and the y-axis the performance (AUROC) of that sub-network.

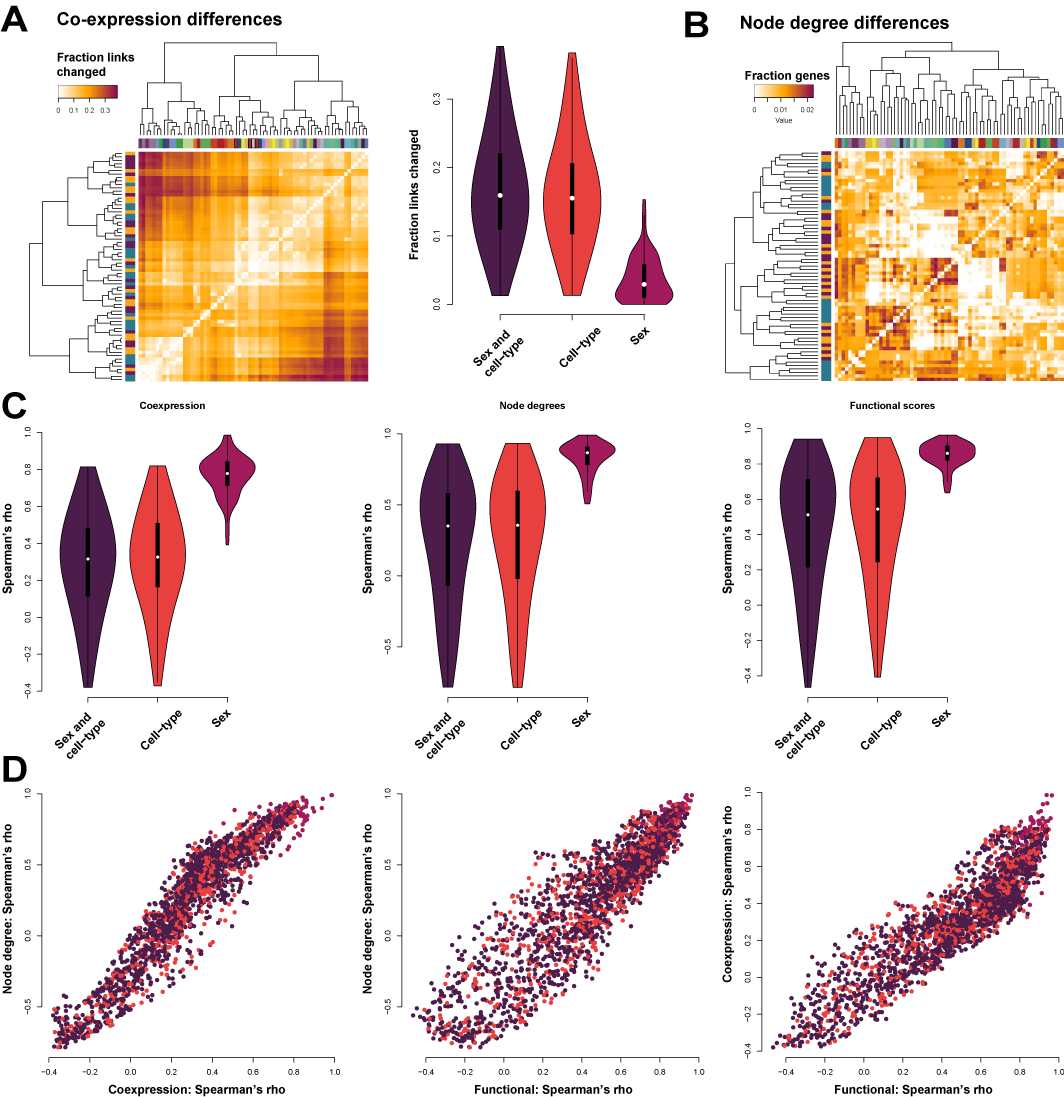

**Supplementary Fig S5 Differential co-expression comparisons.** (A) Proportion of changed links between each aggregate network. (B) Proportion of genes with significantly different node degrees between each aggregate network. (C) Leftpanel: distribution of co-expression correlations between aggregates conditioned on sex and cell-type, cell-type only, or sex only. Middle panel: same as left panel but looking at node degrees and right-panel: network performance AUROCs for GO slim. (D) Comparing correlation values between co-expression, node degree and AUROCs.

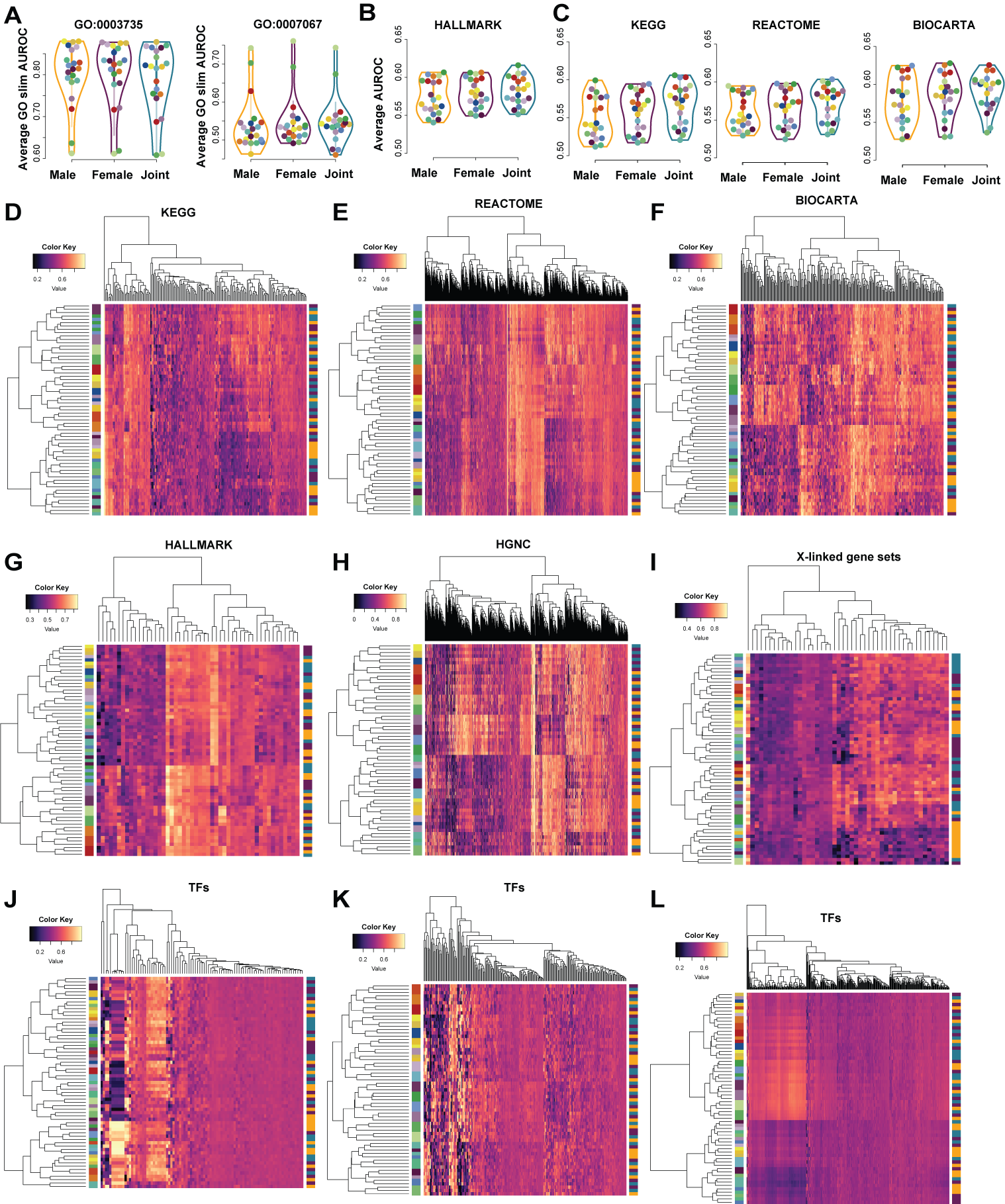

**Supplementary Fig S6 Functional enrichment results of aggregate networks using EGAD.** (A) GO:0003735 structural constituent of ribosome example with high performances for most cell-types. (B) GO:0007067 mitotic cell cycle example with high performances in proliferating cell-types.(C) Average KEGG, REACTOME and BIOCARTA AUROCS. (D) Performance AUROCs for all KEGG, (E) REACTOME, (F) BIOCARTA, (G) HALLMARK, (H) HGNC and (I) curated X-linked gene sets and pathways.
